# Genome-encoded ABCF factors implicated in intrinsic antibiotic resistance in Gram-positive bacteria: VmlR2, Ard1 and CplR

**DOI:** 10.1101/2022.12.05.519065

**Authors:** Nozomu Obana, Hiraku Takada, Caillan Crowe-McAuliffe, Mizuki Iwamoto, Artyom A. Egorov, Kelvin J.Y. Wu, Victoriia Murina, Helge Paternoga, Ben I.C. Tresco, Nobuhiko Nomura, Andrew G. Myers, Gemma C. Atkinson, Daniel N. Wilson, Vasili Hauryliuk

## Abstract

Genome-encoded antibiotic resistance (ARE) ATP-binding cassette (ABC) proteins of the F subfamily (ARE-ABCFs) mediate intrinsic resistance in diverse Gram-positive bacteria. The diversity of chromosomally-encoded ARE-ABCFs is far from being fully experimentally explored. Here we characterise phylogenetically diverse genome-encoded ABCFs from Actinomycetia (Ard1 from *Streptomyces capreolus*, producer of the nucleoside antibiotic A201A), Bacilli (VmlR2 from soil bacterium *Neobacillus vireti*) and Clostridia (CplR from *Clostridium perfringens*, *Clostridium sporogenes* and *Clostridioides difficile*). We demonstrate that Ard1 is a narrow spectrum ARE-ABCF that specifically mediates self-resistance against nucleoside antibiotics. The single-particle cryo-EM structure of a VmlR2-ribosome complex allows us to rationalise the resistance spectrum of this ARE-ABCF that is equipped with an unusually long antibiotic resistance determinant (ARD) subdomain. We show that CplR contributes to intrinsic pleuromutilin, lincosamide and streptogramin A resistance in Clostridioides, and demonstrate that *C. difficile* CplR (CDIF630_02847) synergises with the transposon-encoded 23S ribosomal RNA methyltransferase Erm to grant high levels of antibiotic resistance to the *C. difficile* 630 clinical isolate. Finally, assisted by our novel tool for detection of upstream open reading frames, we dissect the translational attenuation mechanism that controls the induction of *cplR* expression upon an antibiotic challenge.

## INTRODUCTION

Bacterial antimicrobial resistance (AMR) is a constantly growing threat to human health. A 2022 report in *The Lancet* estimated that in 2019 1.27 million deaths were attributable to bacterial AMR globally (Antimicrobial Resistance, 2022). One of the primary antibiotic targets is the ribosome (Arenz and Wilson, 2016), and, accordingly, bacteria protect their protein synthesis machinery via diverse resistance mechanisms (Lin et al., 2018; Wilson, 2014; Wilson et al., 2020). The functional elements primarily targeted by the antibiotics on the large (50S) ribosomal subunit are the peptidyl transferase centre (PTC) and the nascent polypeptide exit tunnel (NPET) (Lin *et al*., 2018; Schwarz et al., 2016). The PTC is targeted by numerous antibiotic classes, including i) nucleoside compounds, such A201A (Polikanov et al., 2015) and hygromycin A (Kaminishi et al., 2015; Polikanov *et al*., 2015), ii) pleuromutilins, such as tiamulin (Schlunzen et al., 2004), iii) lincosamides, such as the natural compound lincomycin (Macleod et al., 1964), its semi-synthetic derivative clindamycin (Phillips, 1981) and the fully synthetic antibiotic iboxamycin (Mitcheltree et al., 2021), iv) type A streptogramins, such as virginiamycin M1 (Tu et al., 2005), v) phenicols, such as chloramphenicol (Syroegin et al., 2022b) and vi) oxazolidinones, such as linezolid (Ippolito et al., 2008; Tsai et al., 2022). Antibiotics targeting the NPET are represented by macrolides, such as the context-dependent polypeptide elongation inhibitor erythromycin (Beckert et al., 2021; Vazquez-Laslop and Mankin, 2018), and by members of the streptogramin B class such as quinupristin (Harms et al., 2004).

The F subfamily of ABC ATPases includes both ribosome-associated antibiotic resistance (ARE) ABCFs (Ero et al., 2019; Wilson *et al*., 2020), as well as multiple housekeeping factors that assist protein synthesis and ribosome assembly (Boël et al., 2014; Chen et al., 2014; Cui et al., 2022; Murina et al., 2019). ARE-ABCF factors mediate antibiotic resistance in human pathogens (exemplified by staphylococcal VgaA (Gentry et al., 2008) and closely related *Listeria monocytogenes* VgaL/Lmo0919 (Crowe-McAuliffe et al., 2021; Dar et al., 2016)), in environmental bacteria (exemplified by *B. subtilis* VmlR (Ohki et al., 2005)), and also confer self-resistance in antibiotic producers (exemplified by *Streptomyces lincolnensis* LmrC (Koberska et al., 2021)). Expression of ARE-ABCFs is suppressed through abortive transcription termination, with full-length mRNA production and ARE-ABCF expression being induced upon antibiotic-induced ribosomal stalling on a regulatory upstream open reading frame (uORF) that is located in the mRNA region preceding the ARE-ABCF main ORF (Cai et al., 2021; Dar *et al*., 2016; Koberska *et al*., 2021; Ohki *et al*., 2005; Takada et al., 2022; Vimberg et al., 2020).

ARE-ABCFs are classified into eight relatively well-studied major subfamilies (ARE1 to ARE8) as well numerous poorly characterised phylum-specific groups, such as actinobacterial AAF1-6 (Crowe-McAuliffe et al., 2022; Murina *et al*., 2019). Members of ARE1-3, ARE5 and ARE6 subfamilies protect the ribosome from PTC-targeting pleuromutilin, lincosamide and streptogramin A (PLS_A_) antibiotics, with the most well-studied ARE-ABCF representatives being ARE1 VgaA and Lmo0919/VgaL, ARE2 VmlR, ARE3 LsaA, ARE5 LmrC, and ARE6 Sal (Crowe-McAuliffe et al., 2018; Dar *et al*., 2016; Novotna and Janata, 2006; Ohki *et al*., 2005; Singh et al., 2002). ARE-ABCFs ARE8 PoxtA and ARE7 OptrA protect against phenicol and oxazolidinone (PhO) transpeptidation inhibitors (Antonelli et al., 2018; Crowe-McAuliffe *et al*., 2022; Wang et al., 2015). Finally, members of the ARE4 subfamily (such as TlrC) and a subset of ARE1 ABCFs (such as Msr) confer resistance to macrolide and streptogramin B (MS_B_) antibiotics (Ross et al., 1990; Rosteck et al., 1991; Su et al., 2018).

ABCF ATPases share a common molecular architecture with two ABC nucleotide-binding domains NBD1 and NBD2 connected by an interdomain linker element which is variable in both amino acid sequence and length (Murina *et al*., 2019). This linker is referred to as the antibiotic resistance determinant (ARD) in the case of ARE-ABCFs (Crowe-McAuliffe *et al*., 2018; Murina et al., 2018; Su *et al*., 2018) and as the P-site tRNA-interaction motif (PtIM) in the case of housekeeping ABCFs (Boël *et al*., 2014; Chen *et al*., 2014). The ARD is crucial for displacement of the antibiotic by ARE-ABCFs, and its deletion renders the resistance determinant inactive (Murina *et al*., 2019). Macrolide-resisting factors such as ARE1 MsrE have the longest ARDs which extend into the ribosomal NPET (Su *et al*., 2018). ARDs of PLS_A_-resisting ARE-ABCF are shorter, as they make contact with the PTC-bound antibiotic and/or induce conformational changes in the PTC that promote antibiotic dissociation (Crowe-McAuliffe *et al*., 2018; Crowe-McAuliffe *et al*., 2021; Mohamad et al., 2022). PhO-resisting PoxtA and OptrA have even shorter ARDs, which act by perturbing the positioning of the CCA-end of the P-site tRNA, rather than directly distorting the PTC (Crowe-McAuliffe *et al*., 2022). Finally, the P-site tRNA-contacting PtIM elements of the housekeeping ABCFs are in general shorter than the ARD elements of ARE-ABCFs (Boël *et al*., 2014; Chen *et al*., 2014; Cui *et al*., 2022).

Antibiotic resistance mediated by ARE-ABCF can synergise with other mechanisms, installed by clinically relevant and widely-spread Erm (Erythromycin resistance methylase) and Cfr (Chloramphenicol and florfenicol resistance) rRNA methyltransferases and confers resistance to macrolide, PhO and PLS_A_ antibiotic classes (Long et al., 2006; Maravić, 2004; Schwarz et al., 2000; Uchiyama and Weisblum, 1985). Recently we have found that *B. subtilis* ARE-ABCF VmlR can synergise with Cfr to grant significant resistance to lincosamides, including the highly potent fully-synthetic antibiotic iboxamycin (Brodiazhenko et al., 2022). The *cfr* gene is often encoded next to *lsa* ABCF in bacterial pathogens such as *Clostridioides difficile* (formerly *Clostridium difficile*) (Gumkowski et al., 2019) and staphylococci (Kehrenberg et al., 2007), which is supportive of the two resistance determinants cooperating to grant high levels of protection. However, synergy of ARE-ABCFs with Cfr/Erm has not as yet been systematically tested experimentally. Importantly, Erm-mediated resistance is associated with an increased risk of *C. difficile* infection (CDI), and the risk further increases upon clindamycin use (Johnson et al., 1999). Neither the role of ARE-ABCFs in *C. difficile* resistance, nor potential ARE-ABCF–Erm synergy, has been studied.

To further understand the molecular mechanisms underlying intrinsic antibiotic resistance in Gram-positive bacteria, we have characterised four genome-encoded ARE-ABCF factors. First, we show that well-studied *B. subtilis* ARE2 VmlR mediates resistance to A201A and hygromycin A, which expands the spectrum of ABCF-mediated resistance to nucleoside antibiotics. Second, we characterise the resistance spectrum of a VmlR representative with an unusually long ARD encoded in the genome of soil bacterium *Neobacillus vireti* (formerly *Bacillus vireti*) – VmlR2. We rationalise the antibiotic sensitivity data by solving the cryo-EM structure of the VmlR2-70S complex and probe its mechanism through mutagenesis. Third, we demonstrate that actinobacterial AAF1 Ard1 encoded by the A201A producer *Streptomyces capreolus* (Barrasa et al., 1995; Saugar et al., 2002) is a narrow spectrum ARE-ABCF that specifically mediates self-resistance against nucleoside antibiotics. Fourth, we establish Clostridial pleuromutilin and lincosamide resistance protein (CplR) as an ARE1 ABCF resistance factor. We characterise CplR of the important human pathogen *C. difficile* (originally annotated as a multidrug resistance (MDR)-type ABC transporter CDIF630_02847 (Fuchs et al., 2021)), of *Clostridium perfringens*, a common causative agent of food poisoning, as well as of the mutualistic bacterium *Clostridium sporogenes*. We demonstrate that CplR contributes to well as pleuromutilin retapamulin. While lincosamide resistance mediated by CplR can be overcome by the fully synthetic lincosamide iboxamycin (Mitcheltree *et al*., 2021), ARE1 CplR provides high levels of iboxamycin resistance when acting together with transposon-encoded ErmB present in clinically isolated *C. difficile* strain 630 (Farrow et al., 2001). Given that clindamycin treatment is associated with increased risk of CDI (Bishara et al., 2008; Buffie et al., 2012; Duffy et al., 2020; Johnson *et al*., 1999; Sullivan et al., 2001), our discovery and characterisation of CplR-mediated lincosamide resistance mechanism in *C. difficile* is of direct clinical importance.

## MATERIALS AND METHODS

### Sequence and structure analysis

dRNA-seq and RNAtag-seq data signal tracks (Fuchs *et al*., 2021) were downloaded from the NCBI GEO (GSE155167) and visualised using svist4get (Egorov et al., 2019). Searching for conserved short ORFs in 5ʹ leader region sequences of *cplR*, *vmlR* and *vmlR2* was performed with uORF4u [https://github.com/art-egorov/uorf4u] (Egorov and Atkinson, 2022). The pipeline consists of several steps: i) BlastP for homology searching (Camacho et al., 2009) with cut-offs set on a protein by protein basis to optimise numbers and diversity of hits: 50% sequence identity to the query cutoff for CplR (WP_011861613.1), 70% identity cutoff for VmlR (WP_003234144.1) and VmlR2 (WP_024026878.1) and 80% for LsaA (WP_002398829.1), ii) retrieval of 300 nt upstream sequences preceding the identified ARE-ABCF genes, iii) uORF annotation and, finally, iv) conservation analysis and multiple sequence alignment with MAFFT v7.490 (Katoh and Standley, 2013). Visualisation was performed with the ggmsa R package (Zhou et al., 2022), msa4u (Egorov and Atkinson, 2022) and Logomaker (Tareen and Kinney, 2020) Python packages. Secondary structures of the *cplR* 5ʹ leader region were predicted using RNAfold (Gruber et al., 2008) and Mfold (Peltier et al., 2020). For phylogenetic analysis, selected ARE-ABCFs were aligned with MAFFT v7.490 with the L-INS-i strategy (Katoh and Standley, 2013), and visualized with AliView v1.26 (Larsson, 2014) and Jalview v2.11.2.0 (Waterhouse et al., 2009). Alignment positions with >50% gaps were removed with trimAI v1.4.rev6 (Capella-Gutierrez et al., 2009) before tree building with IQTree v2.1.2 on the CIPRES server with 1000 rapid bootstrap replicates and the model ‘LG-I-G4’, as selected using the automatic model detection setting (Miller et al., 2010; Minh et al., 2020).

### Bacterial culture

*C. difficile* 630Δ*erm* and its derivatives were grown anaerobically in Brain-Heart Infusion (Difco) supplemented with 0.5% yeast extract and 0.1% cysteine (BHIS) or BHIS agar under 80% N_2_, 20% CO_2_, 4% H_2_ atmosphere in the Coy anaerobic chamber (COY). *C. sporogenes* and its derivatives were anaerobically grown in Gifu anaerobic medium (GAM) (Nissui Co., Japan) in the Coy anaerobic chamber. *C. perfringens* and its derivatives were grown at 37 °C in GAM under anaerobic conditions using Anaeropack system (Mitsubishi Gas Chemical Co. Inc., Tokyo, Japan). *B. subtilis* were routinely aerobically cultured in LB broth. When necessary, antibiotics were supplemented in the media: 20 µg/mL erythromycin and 10 µg/mL thiamphenicol in the case of *C. difficile* and 20 µg/mL chloramphenicol in the case of *C. perfringens*.

### Strain and plasmid construction

The plasmid and oligonucleotides used in this study are listed in **Supplementary Table 1**, respectively.

To construct *C. difficile cplR* (CDIF630_02847) deletion mutant we constructed the plasmid for the allele exchange mutagenesis as previously reported by Peltier and colleagues (Peltier *et al*., 2020). The CD2517.1 type I toxin gene under the control of the tetracycline-inducible promoter as well as ≈1 kb-long 5′ and 3′ flanking regions of the target gene were cloned into pMTL83151 plasmid (CHAIN biotech, Nottingham, UK), yielding the pMSRNO plasmid. Using *E. coli* HB101/pRK24, the pMSRNO plasmid was then introduced into *C. difficile* by conjugation. pRK24 (Ma et al., 2014) was a gift from Farren Isaacs (Addgene plasmid # 51950). Briefly, 500 µL of overnight cultured *C. difficile* were heated at 52 °C for 2 minutes. Meanwhile, 1 mL culture of *E. coli* HB101/pRK24 harbouring pNO201 were centrifuged and pelleted at 1500 × *g* for 3 minutes. The *E. coli* pellet was suspended to 200 µl of the heated *C. difficile* culture in the anaerobic chamber. The mixture was spotted onto a BHI agar plate and incubated for 8 hours. The mixed colonies were resuspended in 1 mL BHI broth, and then spread onto BHIS plates supplemented with 10 µg/mL thiamphenicol, 250 µg/mL D-cycloserine and 50 µg/mL kanamycin (BHIS-TCK). After incubation for two to three days, individual colonies were restreaked on BHIS-TCK for isolation. The protocol for allele exchange was based on that of Peltier and colleagues (Peltier *et al*., 2020). The deletion of the target gene was confirmed by PCR amplification.

*C. sporogenes* mutants were constructed using the Targetron system (Merck). A DNA fragment containing group II intron and *ltrA* gene from pJIR750ai (Merck) as well as a *Clostridium-E. coli* shuttle vector pMTL83153 (CHAIN biotech, Nottingham, UK) containing a constitutive promoter P*_fdx_* were PCR-amplified, and the intron-containing plasmid was produced by ligation of these DNA fragments pre-digested with *Hind*III and *Xho*I. Then, a PCR-amplified *C. sporogenes cplR*-targeted intron was introduced into *Hind*III-*BsrG*I site in the plasmid. The resulting pNO201 plasmid was used for *cplR* mutant construction. The pNO201 plasmid was introduced into *C. sporogenes* by conjugation as described above, with one modification: the heat treatment prior to conjugation was omitted. While this step is crucial for obtaining transconjugants with *C. difficile*, in the case of *C. sporogenes* transconjugants are easily obtained without heat shock. To screen the mutant in which the target gene was disrupted, the colonies were restreaked onto BHIS containing 20 µg/mL erythromycin. The insertion of the intron in the appropriate locus was verified by PCR amplification.

The *C. perfringens* mutant was constructed as described previously (Nariya et al., 2011; Obana et al., 2020). Briefly, ≈1 kb of the 5′ and 3′ flanking regions of the target gene were PCR-amplified, and the resultant fragments were cloned into pCM-GALK using the In-Fusion cloning system (Takara Bio, Japan). Resulting plasmid was introduced into *C. perfringens* HN13 via electroporation, and the desired mutant was isolated on agar plates supplemented with 20 μg/mL chloramphenicol or 3% galactose. The deletion of the target gene was confirmed by PCR and DNA sequencing.

For complementation of *cplR* genes in *C. difficile* and *C. perfringens*, *cplR*-expressing plasmids were constructed. The *C. difficile cplR*-expressing plasmid was based on pRPF185 plasmid which contains a tetracycline-inducible promoter (Fagan and Fairweather, 2011). The pRPF185 was a gift from Robert Fagan and Neil Fairweather (Addgene plasmid # 106367; http://n2t.net/addgene:106367; RRID: Addgene_106367). A PCR-amplified *cplR* CDS was cloned into pRFP185 using the In-Fusion cloning system. For complementation of *C. perfringens cplR*, the DNA fragment containing a native promoter and CDS of *C. perfringens cplR* gene was amplified by PCR and cloned into pJIR750 (Bannam and Rood, 1993).

An *mCherry2-L* variant codon-optimized for low-GC bacteria was used to construct the reporter constructs used to assay the regulation of *cplR* expression (Ransom et al., 2015). The DNA fragment encoding a codon-optimized *mCherry2-L* CDS was synthesized at Eurofin Genomics (Tokyo, Japan). DNA fragments containing *cplR* promoter and 5′ leader region were amplified and ligated with the *mCherry2-L* by PCR. The resulting DNA was introduced into pMTL84151 (CHAIN biotech) using the In-Fusion cloning system.

The *B. subtilis* strains were derivatives of wt168 (*trpC2*). The strains were constructed through homologous recombination by transformation with plasmids listed in **Supplementary Table 1**. The plasmids were constructed by standard cloning methods including PCR and Gibson assembly using oligonucleotides listed in **Supplementary Table 1**. Successful integration of a gene into the chromosome was accomplished by double crossing-over at the target loci. The resulting recombinant clones were checked for their antibiotic resistance markers, including the absence of those originally present on the plasmid backbone, as well as for inactivation of the *thrC* target locus.

### Antibiotic susceptibility testing

The Minimum Inhibitory Concentrations (MIC) were calculated according to guidelines from the European Committee on Antimicrobial Susceptibility Testing (EUCAST) (http://www.eucast.org/ast_of_bacteria/mic_determination). Bacterial cells were cultured in 96-well plates in medium supplemented with 2-fold serial dilutions of the antibiotics. BHIS, GAM, or LB media was used to grow *C. difficile*, *C. perfringens*/*C. sporogenes* and *B. subtilis*, respectively. The 200 µL cultures were inoculated at the concentration of 5 × 10^5^ CFU/mL and growth in the 96-well plates. The MIC was determined as the lowest concentration of antibiotics in which no bacterial growth was observed after incubation for 24 hours at 37 °C. The reproducibility was confirmed by at least 3 individual experiments.

### Immunoprecipitation of C-terminally HTF-tagged VmlR2_N.vireti_^EQ2^

The experiments were reformed as described earlier (Crowe-McAuliffe *et al*., 2021). C-terminally HTF-tagged VmlR2*_N.vireti_*^EQ2^ inducible strain VHB220 (Δ*vmlR thrC*::P*_hy-spnak_*-*vmlR_B.vireti_*^EQ2^*-HTF*; HTF stands for His_6_-TEV-FLAG_3_) was pre-grown on LB plates overnight at 30 °C. Fresh individual colonies were used for inoculation and grown in LB medium with 1 mM IPTG. 0.5 L at –80°C. Cell pellets were resuspended in 1 mL of cell opening buffer (95 mM KCl, 5 mM NH_4_Cl, 20 mM HEPES (pH = 7.5), 1 mM DTT, 15 mM Mg(OAc)_2_, 0.5 mM CaCl_2_, 8 mM putrescine, 1 mM spermidine, 1 tablet of cOmplete™ EDTA-free Protease Inhibitor Cocktail (Roche) per 50 mL of buffer) and disrupted using FastPrep homogeniser (MP Biomedicals) with 0.1 mm Zirconium beads (Techtum) in 6 cycles by 20 seconds with 3-minute chilling on ice. Cell debris was removed by centrifugation at 14,800 rpm for 20 minutes 4 °C in F241.5P rotor using 149 Microfuge 22R centrifuge (Beckman Coulter). The supernatant was combined with 100 µL of ANTI-FLAG M2 Affinity Gel (Sigma) pre-equilibrated in cell opening buffer, and incubated for 1.5 hours at 4 °C on a turning wheel (Fisherbrand™ Multi-Purpose Tube Rotators). The samples were loaded on Micro Bio-Spin columns (Bio-Rad) pre-equilibrated in cell opening buffer, and washed 5 times with 0.5 mL of cell opening buffer by gravity flow. VmlR2*_vireti_*^EQ2^-HTF was eluted by addition of 200 µL opening buffer containing 0.1 mg/mL poly-FLAG peptide (Biotool, Bimake) for 45 min on a turning wheel. All incubations, washes and elutions were performed at 4 °C. The eluted sample was collected by centrifugation at 2000 rpm for 1 minute 4 °C in a F241.5P rotor using a 149 Microfuge 22R centrifuge (Beckman Coulter). For SDS-PAGE analyses, 20 µL aliquots of samples were mixed with 5 µL of 5x SDS loading buffer and heated at 95 °C for 15 minutes, and denatured samples were loaded on 12% SDS-PAGE. SDS-gels were stained by “Blue-Silver” Coomassie Staining (Candiano et al., 2004) and washed with water for 6 hours or overnight before imaging with LAS4000 (GE Healthcare).

### Preparation of cryo-EM grids

Elutions from pull-downs were kept on ice until being applied within two hours to glow discharged cryo-grids (Quantifoil 2/2 Cu_300_ coated with 2 nm continuous carbon). 3.5 µL of sample was loaded on grids in Vitrobot (FEI) under conditions of 100% humidity at 4 °C, blotted for 5 seconds and vitrified by plunge-freezing in liquid ethane. Samples were imaged on a Titan Krios (FEI) operated at 300 kV at a nominal magnification of 165,000 x (0.82 Å/pixel) with a Gatan K2 Summit camera and a BioQuantum energy filter with a slit width of 20 eV. The exposure rate was 5.277 electrons/pixel/s with a 4 second exposure and 20 frames using the EPU software.

### Cryo-EM data analysis

MotionCor2 was used to correct for beam-induced motion in 6,384 starting micrographs (Zheng et al., 2017). Contrast transfer functions were estimated with Gctf (Zhang, 2016). Subsequent processing was performed in RELION 3.1 unless specified otherwise (Scheres, 2012; Zivanov et al., 2018). Micrographs with MaxRes > 4 Å and/or a CtfFigureOfMerit < 0.05 were discarded, resulting in 5,014 micrographs used for further processing. From the resulting set of 5,014 micrographs, 279,082 particles were picked with Gautomatch [https://www2.mrc-lmb.cam.ac.uk/download/gautomatch-056] and extracted with a pixel size of 2.46 Å. 2D classification was used to remove non-ribosomal particles, resulting in 193,343 particles which were subjected to 3D auto refinement. The initial reference, which was low-pass filtered to 60 Å, was a *B. subtilis* 70S ribosome that contained a P-tRNA and no factor in the E site (EMD-0176, (Crowe-McAuliffe *et al*., 2018)). The output volume was used as a reference for subsequent 3D refinements. 3D classification with four classes and without angular sampling was then performed, resulting in two classes of interest that contained density in the E site. The best-resolved of the two classes, class 2, was used for most analyses, on the grounds that this class had the most continuous and feature-rich density for the protein in the E site. Particles from the class 2 were re-extracted at the original pixel size and used for 3D autorefinement, and then anisotropic magnification, per-particle defocus, per-micrograph astigmatism, beam tilt, and higher-order aberrations were refined before final 3D autorefinement and post-processing (Zivanov *et al*., 2018).

A combined volume, containing particles from both classes with VmlR density in the E site, was also re-extracted as above and partial signal subtraction followed by 3D classification around the A and E sites was performed. For the A-site classification, three classes with 29.1%, 37.6%, and 33.3% occupancy were observed. The first and last of these classes contained a tRNA, each in a slightly different conformation, while the second class had no density in the A site, consistent with the poorly resolved density for the A tRNA in the parent map. Each of these classes was reverted to full density and used for 3D autorefinement. Partial signal subtraction and 3D classification around the E site and L1 stalk yielded three maps that differed mostly in the conformation of the L1 stalk, consistent with the poor resolution of this region in the parent map. However, density for VmlR2 was not improved in any of these classes, so class 2 from the initial 3D classification was used for further analyses.

For post-processing, soft solvent masks and a B factor automatically estimated by Guinier analysis were used. See also **Supplementary Figure 3** for an overview of processing and **Supplementary Table 2** for collection and processing statistics.

### Model building and refinement

Starting models for *B. subtilis* ribosomal proteins and VmlR2 were obtained from the AlphaFold2 database (Varadi et al., 2022). Ribosomal RNA was taken from PDB 6HA1 (Crowe-McAuliffe *et al*., 2018). Models were first adjusted using Coot (Casanal et al., 2020) into a high-resolution *B. subtilis* 70S map (to be described elsewhere). The resulting model was then fitted in the VmlR2-70S map and adjusted manually with Coot. The starting model of VmlR2 was fitted domain-by-domain and adjusted by hand. A prediction created by ModelAngelo was also used to guide modelling of VmlR2 in places (Jamali et al., 2022). For the P-site tRNA and 23S rRNA of the L1 stalk, PDBs 7NHK and 7NHL were used as starting points, respectively (Crowe-McAuliffe *et al*., 2021). Density for the uL1 protein and the CCA-3′ end nucleotides of the P-site tRNA was very unclear and these regions were therefore omitted from the model. PDB 7K00 was used as a reference to guide modelling, especially around 23S rRNA helix 69 (Watson et al., 2020). The resulting model was refined first with REFMAC5/Servalcat (Kovalevskiy et al., 2018; Yamashita et al., 2021), then with Phenix using the starting model as a reference for restraints to decrease bond length and angle root-mean-square deviations (Liebschner et al., 2019). MolProbity through Phenix was used for model validation (Williams et al., 2018). Model statistics are available in **Supplementary Table 2**. Figures showing maps and models were made with UCSF ChimeraX (Pettersen et al., 2021) or PyMOL (https://github.com/schrodinger/pymol-open-source).

### RNA extraction and RT-PCR

Overnight culture 630Δ*erm C. difficile* strain in BHIS was diluted 1:20 in fresh BHIS medium and incubated anaerobically for 2 hours to reach the log phase. Then we added various concentrations of antibiotics to the culture. After incubation for 0.5 to 6 hours, the cells were collected from 1 mL of the culture by centrifugation at 10,000 × *g* for 2 minutes and flash frozen in liquid nitrogen. Total RNAs were extracted as described previously (Obana et al., 2010) with minor modifications. Briefly, the cell pellets were flash-frozen in liquid nitrogen until use. The phenol/chloroform/isoamylalchol followed by shaking for 5 minutes using a Shake Master NEO (Bio Medical Science, Tokyo, Japan). After centrifugation at 20,000 × *g* for 5 minutes, the aqueous layer was transferred to a fresh tube and mixed with equal volume of phenol/chloroform/isoamylalchol. To recover the total RNAs, the aqueous layer was mixed with 2.5 volumes of ethanol and 0.1 volumes of 3 M NaOAc, pH5.2. The RNA was harvested by centrifugation at 20,000 × *g* for 10 minutes, washed with 70% ethanol, and contaminated DNA was removed by DNase I treatment (Promega). Purified RNA samples were resuspended in 50 μL of RNase-free water and stored at −80 °C.

Reverse transcription was preformed with PrimeScript™ RT Master Mix kit (TaKaRa, Japan) using 10 ng or 0.5 ng (genomic or plasmid encoding genes, respectively) of total RNA. The cDNAs for the target genes were amplified using Tks Gflex™ DNA Polymerase (TaKaRa, Japan) and the primers used are listed in **Supplementary Table 1**.

### mCherry fluorescent reporter assays

*C. difficile* harbouring reporter plasmids were cultured in BHIS supplemented with 10 μg/mL thiamphenicol. Cells were harvested from 1 mL cultures by centrifugation at 10,000 × *g* for 2 min and washed with phosphate buffer saline (PBS). The resultant cell pellets were fixed with 4% formaldehyde for 8 minutes, washed with PBS and resuspended in 1 ml of PBS. After exposure to oxygen for 90 minutes for chromophore maturation, fluorescence (ex/em = 579/616 nm) and absorbance (600 nm) of each sample were measured using a Synergy H1 microplate reader (BioTek, Vermont, USA). To calculate the relative promoter activity, relative fluorescence intensity normalised by O.D._600_ was further normalised by the signal from the constitutive P*_fdx_-mCherry* reporter.

## RESULTS

### The ABCF ARE1 subfamily member CplR is conserved in Clostridioides

While horizontally mobile ARE-ABCFs that are encoded in mobile genetic elements or plasmids mediate acquired antibiotic resistance, chromosomally-encoded ARE-ABCFs provide intrinsic antibiotic resistance and are likely to be predominantly inherited vertically. The three Gram-<u>p</u>leuromutilin and lincosamide resistance; we propose this new name based on the resistance spectrum determined in the current study, see below), VmlR2 and Ard1 – are of the latter, intrinsic type. Despite this commonality, they belong to very different subfamilies of the ABCF family tree (**Figure 1A**). *C. difficile* CplR encoded by the gene CDIF630_02847 (RefSeq protein accession WP_011861613.1) is a member of the ARE1 subfamily, although it is phylogenetically distinct from the well-characterised mobile Vga- and Msr-like ARE1 proteins. CplR is found in a range of different Clostridia, including *C. difficile*, *C. perfringens* and *C. sporogenes* (**Figure 1A**). The ARE2 subfamily of ABCFs can be subdivided into two phylogenetically distinct types: one subtype has a short ‘arm’ subdomain, as is seen in VmlR from *B. subtilis* (Crowe-McAuliffe *et al*., 2018), while the second form has insertions in the arm and – sometimes – the ARD/linker element, as is the case with *Neobacillus vireti* VmlR2 (**Figure 1B**, **Supplementary Figure 1**). The Ard1 ABCF from *Streptomyces capreolus* is classified as being in the actinobacterial subfamily AAF1, closely related to ARE4 and ARE5 subfamilies (Murina *et al*., 2019). Just as with *C. difficile* CplR ARE-ABCF (Fuchs *et al*., 2021), Ard1 was originally mis-annotated as a transporter (Barrasa *et al*., 1995). While ABCF proteins do carry the ATP-binding cassette domain that is also found in many transmembrane transporters, they lack the necessary transmembrane domains for such a role. Ard1 is encoded as part of the A201A antibiotic biosynthesis gene cluster, acting as a self-resistance determinant (Saugar *et al*., 2002).

**Figure 1.**
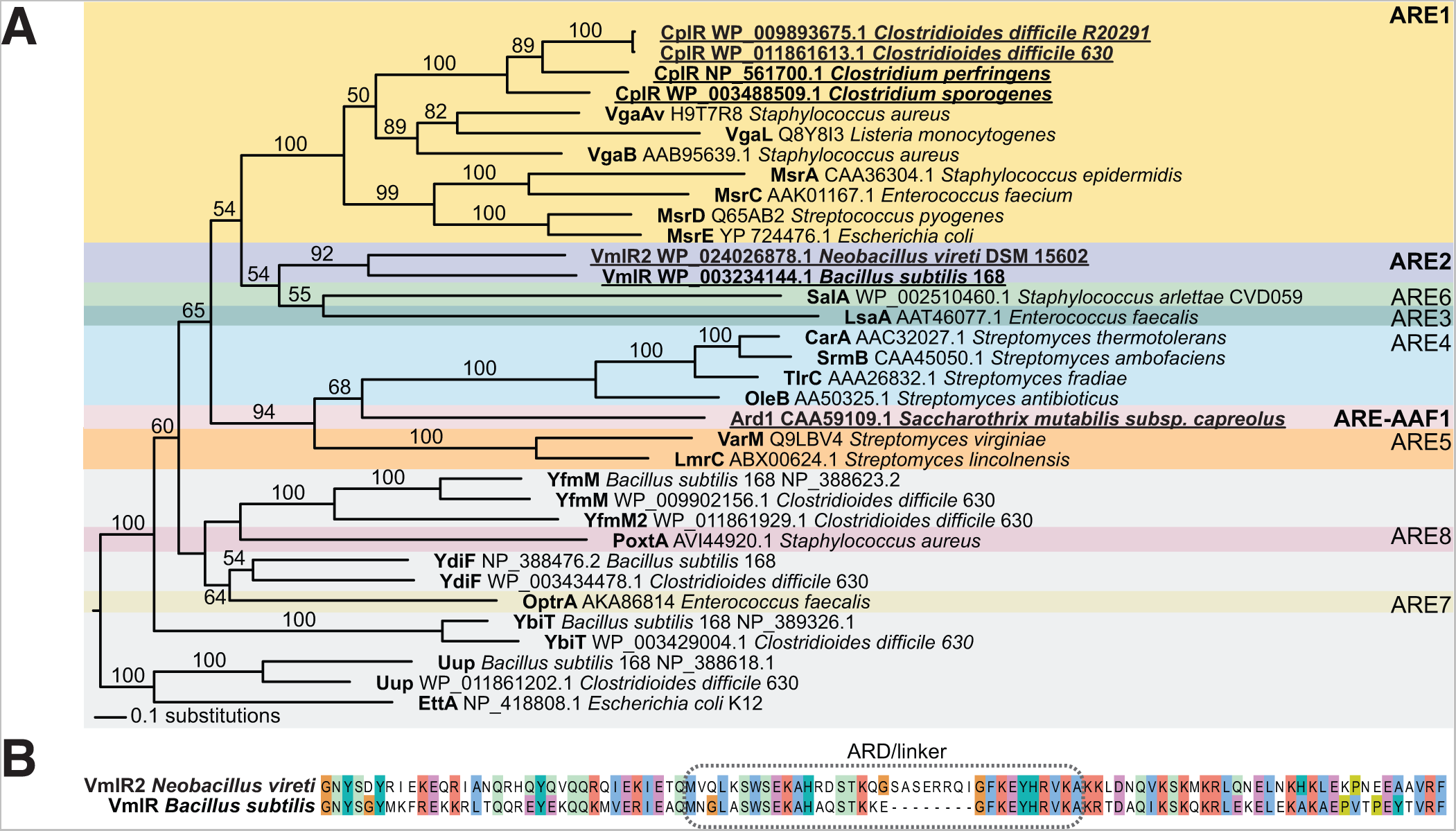
Maximum likelihood phylogenetic analysis of selected ABCFs places CplR, Ard1 and VmlR2 in different clades of the ABCF family tree. **A**) Branches are proportional to the number of amino acid substitutions according to the lower left legend. Branch support values are percentage Ultrafast Bootstrap Support values, in percentages. Only those 50% or higher are shown. CplR is a member of the ARE1 subfamily, but without any strongly supported specific association with Vga or Msr subtypes. There is strong support for Ard1 falling within the same clade as other actinomycete ARE subfamilies, all of which likely confer self-protection of the antibiotic producer. VmlR2 is a close relative of VmlR, but has distinctive sequence/structure features, including (**B**) an extended ARD/linker subdomain (see **Supplementary Figure 1** for full alignment).

### Antibiotic resistance spectra of *S. capreolus* AAF1 Ard1, *N. vireti* ARE2 VmlR2, *C. difficile* ARE1 CplR expressed in *B. subtilis* Δ*vmlR* surrogate host

To determine the spectra of antibiotic resistance conferred by *N. vireti* VmlR, *S. capreolus* Ard1, and *C. difficile* CplR, we first ectopically expressed these ARE-ABCFs in a *B. subtilis* surrogate host under the control of an IPTG-inducible P*_hy-spank_* promoter (Britton et al., 2002) and determined the Minimum Inhibitory Concentrations (MICs) for a comprehensive panel of translation-targeting antibiotics (**Table 1**). We used a *B. subtilis* strain lacking the VmlR ARE-ABCF factor since the Δ*vmlR* strain is highly sensitive to PLS_A_ antibiotics (Brodiazhenko *et al*., 2022; Crowe-McAuliffe *et al*., 2018; Ohki *et al*., 2005; Takada *et al*., 2022).

**Table 1.** Resistance profiles of VmlR, VmlR2, Ard1 and CplR ARE-ABCFs expressed in B. subtilis ΔvmlR surrogate host. B. subtilis MIC testing for a panel of nucleoside, PLS_A_ and phenicol antibiotics was carried out in either LB medium or LB supplemented with 1 mM IPTG (to induce expression of ARE-ABCF proteins), and growth inhibition was scored after 16-20 hours at 37 °C. N.D. stands for ‘not determined’.

| *B. subtilis* strain | MIC (µg/ml) |  |  |  |  |  |  |  |  |
| --- | --- | --- | --- | --- | --- | --- | --- | --- | --- |
|  | A201A | Hygromycin A | Retapamulin | Tiamulin | Lincomycin | Iboxamycin | Virginia mycin M1 | Chloramphenicol | Erythromycin |
| 168 wt | >256 | >256 | 80 | 80 | 80 | 2 | >64 | 5 | 0.125 |
| $\Delta vmlR$ | 4 | 16 | 0.078 | 0.3 - 0.6 | 2.5 | 0.06 | 2 | 5 | 0.125 |
| $\Delta vmlR$ thrC::P<sub>hy-spank</sub>-vmlR (IPTG: 1 mM) | >256 | >256 | >80 | >80 | >80 | >2 | >64 | 5 | 0.125 |
| $\Delta vmlR$ thrC::P<sub>hy-spank</sub>-vmlR<sup>EQ2</sup> (IPTG: 1 mM) | 2-4 | 16 | 0.078 | 0.3-0.6 | 2.5 | 0.06 | 2 | 5 | 0.125 |
| $\Delta vmlR$ thrC::P<sub>hy-spank</sub>-vmlR2<sup>N.vireti</sup> (IPTG: 1 mM) | >256 | >256 | >80 | >80 | >80 | 8 | >64 | 5 | 0.125 |
| $\Delta vmlR$ thrC::P<sub>hy-spank</sub>-vmlR2<sup>N.vireti</sup>-HTF (IPTG: 1 mM) | N.D. | N.D. | >80 | >80 | >80 | 8 | >64 | 5 | 0.125 |
| $\Delta vmlR$ thrC::P<sub>hy-spank</sub>-vmlR2<sup>N.vireti</sup><sup>EQ2</sup>-HTF (IPTG: 1 mM) | N.D. | N.D. | 0.078 | 0.3-0.6 | 2.5 | 0.03 | 2 | 5 | 0.125 |
| $\Delta vmlR$ thrC::P<sub>hy-spank</sub>-vmlR2<sup>N.vireti</sup><sup>R259A</sup> (IPTG: 1 mM) | N.D. | N.D. | 80 | N.D. | >80 | 2 | >64 | 5 | N.D. |
| $\Delta vmlR$ thrC::P<sub>hy-spank</sub>-vmlR2<sup>N.vireti</sup><sup>Q260A</sup> (IPTG: 1 mM) | N.D. | N.D. | >80 | N.D. | >80 | 8 | >64 | 5 | N.D. |
| $\Delta vmlR$ thrC::P<sub>hy-spank</sub>-vmlR2<sup>N.vireti</sup><sup>I261A</sup> (IPTG: 1 mM) | N.D. | N.D. | >80 | N.D. | >80 | 8 | >64 | 5 | N.D. |
| $\Delta vmlR$ thrC::P<sub>hy-spank</sub>-vmlR2<sup>N.vireti</sup><sup>E265A</sup> (IPTG: 1 mM) | N.D. | N.D. | 1.25> | N.D. | 80 | 0.25 | 16 | 5 | N.D. |
| $\Delta vmlR$ thrC::P<sub>hy-spank</sub>-vmlR2<sup>N.vireti</sup><sup>Y266A</sup> (IPTG: 1 mM) | N.D. | N.D. | 2.5 | N.D. | >80 | 2 | >64 | 5 | N.D. |
| $\Delta vmlR$ thrC::P<sub>hy-spank</sub>-ard1 (IPTG: 1 mM) | 16 | 32 | 0.078 | 0.3-0.6 | 2.5 | N.D. | 2 | 5 | 0.125 |
| $\Delta vmIR$ $P_{hy-spank^-}$<br>$ard1^{EQ2}$ (IPTG: 1 mM) | 4 | 16 | 0.078 | 0.3-0.6 | 2.5 | N.D. | 2 | 5 | 0.125 |
| $\Delta vmIR$ $P_{hy-spank^-} cplR^{Cd}$<br>(IPTG: 1 mM) | 16 | N.D. | >80 | >80 | >80 | 1-2 | >64 | 5 | 0.125 |
| $\Delta vmIR$ $P_{hy-spank^-}$<br>$cplR^{Cd}$-FLAGx3<br>(IPTG: 1 mM) | 4 | N.D. | 0.078 | 0.3-0.6 | 2.5 | N.D. | 2 | 5 | N.D. |

While the wild-type *B. subtilis* is virtually immune to A201A and hygromycin A (MICs in excess of 256 µg/mL, beyond the concentration levels that are experimentally feasible given the paucity of these antibiotics), the Δ*vmlR* strain is sensitive (MIC for A201A of 4 µg/mL, MIC for hygromycin A of 16 µg/mL) (**Table 1**). This observation expands the spectrum of known VmlR-mediated resistance to nucleoside compounds. When *B. subtilis* VmlR is overexpressed in Δ*vmlR B. subtilis*, the strain becomes PLS_A_- and nucleoside-resistant; no protective effect is detectable when an ATPase-deficient EQ_2_ variant of VmlR is expressed (**Table 1**). Expression of *N. vireti* VmlR2 confers the same PLS_A_ and nucleoside resistance spectrum, despite the relatively long ARD, without any detectable resistance to macrolides (**Table 1**). The P*_hy-spank_*- driven expression of wild-type *S. capreolus* Ard1 – but not that of EQ_2_ Ard1 – confers Δ*vmlR B. subtilis* moderate resistance to nucleoside – but not PLS_A_ – antibiotics. The narrow spectrum of Ard1-mediated resistance suggests that Ard1 specifically confers self-resistance to A201A (Murina *et al*., 2019) (**Table 1**). Finally, ectopic expression of *C. difficile* CplR confers Δ*vmlR B. subtilis* resistance to PLS_A_ antibiotics and has a moderate (2- to 4-fold increase in MIC) protective effect against nucleosides.

Collectively, these results i) expand the resistance spectra of ABCF-mediated resistance to nucleoside antibiotics, ii) demonstrate that despite its relatively long ARD, *N. vireti* ARE2 VmlR has a PLS_A_ and nucleoside resistance spectrum identical to that of the short-ARD *B. subtilis* ARE2 VmlR, iii) establish *S. capreolus* Ard1 ARE-AAF1 as a dedicated narrow-spectrum nucleoside resistance factor and iv) establish genome-encoded *C. difficile* ARE1 CplR as a PLS_A_ resistance determinant.

### Cryo-EM structure of *N. vireti* VmlR2-70S complex

Next, we aimed to determine cryo-electron microscopy structures of ribosome-bound ATPase-deficient (EQ_2_) *C. difficile* ARE1 CplR and *N. vireti* ARE2 VmlR2. To generate the ribosome-bound complexes we employed our well-established affinity purification strategy (Crowe-McAuliffe *et al*., 2022; Crowe-McAuliffe *et al*., 2021). While the *C. difficile* CplR structure would provide the necessary structural insights into intrinsic PLS_A_ and nucleoside antibiotic resistance of *C. difficile* pathogen, the *N. vireti* VmlR structure would be instructive for structural rationalisation of the VmlR2*_N.vireti_* resistance spectrum that is identical to that of short-ARD *B. subtilis* VmlR. *S. capreolus* Ard1 has not been prioritised for structural studies for two reasons: i) the lack of established genetic tools for *S. capreolus* and ii) the levels of nucleoside antibiotic resistance conferred in *B. subtilis* surrogate host are modest, indicative of imperfect functioning of the resistance determinant on the *B. subtilis* 70S ribosome.

Unfortunately, our attempts to purify the FLAG_3_-tagged *C. difficile* CplR in complex with the ribosome were unsuccessful. As C-terminal FLAG_3_-tagging of *C. difficile* CplR abrogated the functionality of CplR in Δ*vmlR B. subtilis* (**Table 1**; Δ*vmlR thrC*::P*_hyspank_*-*cplR^Cd^-FLAG_3_*), we did not pursue this direction further. However, we were successful in isolating a complex of EQ_2_ *N. vireti* VmlR2-HTF with *B. subtilis* 70S ribosomes (**Supplementary Figure 2**). The immunoprecipitated FLAG_3_-tagged EQ_2_ VmlR2 was eluted and applied to a cryo-EM grid for imaging in a Titan Krios followed by single-particle analysis using RELION (Scheres, 2012). Three-dimensional classification revealed one major class of particles (72.3% of particles after 2D classification) consisting of VmlR2-EQ_2_ bound to the 70S ribosome (**Supplementary Figure 3**). This volume could be refined to an average resolution of 2.9 Å (**Supplementary Figure 4A,B**). The VmlR2 NBDs and C-terminal extension (CTE) could be modelled by domain-wise fitting and adjustment of the AlphaFold2 model (**Supplementary Figure 4C**). While the density for the NBDs was not sufficient to draw conclusions about the precise geometry adopted by the catalytic region of the enzyme, the VmlR2 interdomain linker, consisting of two α helices and a loop, and which extends into the core of the large ribosomal subunit, was well resolved and could be modelled *de novo* with high confidence (**Supplementary Figure 4C**). The VmlR2-70S structure is globally similar to other antibiotic resistance ABCF-70S complexes (Crowe-McAuliffe *et al*., 2018; Crowe-McAuliffe *et al*., 2022; Crowe-McAuliffe *et al*., 2021; Mohamad *et al*., 2022; Su *et al*., 2018) (**Figure 2A,B**). VmlR2-EQ_2_ is bound in the ribosomal E site with closed NBDs and density corresponding to two bound NTP molecules, presumably ATP (**Figure 2A-C**). The VmlR2 α-helical interdomain linker extends from the E site into the PTC, overlapping the canonical position of a P-site tRNA (**Figure 2D,E**). This results in a distorted tRNA occupying the P site, with the tRNA elbow contacting the VmlR2 NBD2, pulled towards the E site, and the acceptor stem highly distorted. Poor density for the P-tRNA acceptor stem indicates a high degree of flexibility in this region, and in particular the CCA-3′ end could not be modelled (**Supplementary Figure 5A**).

**Figure 2.**
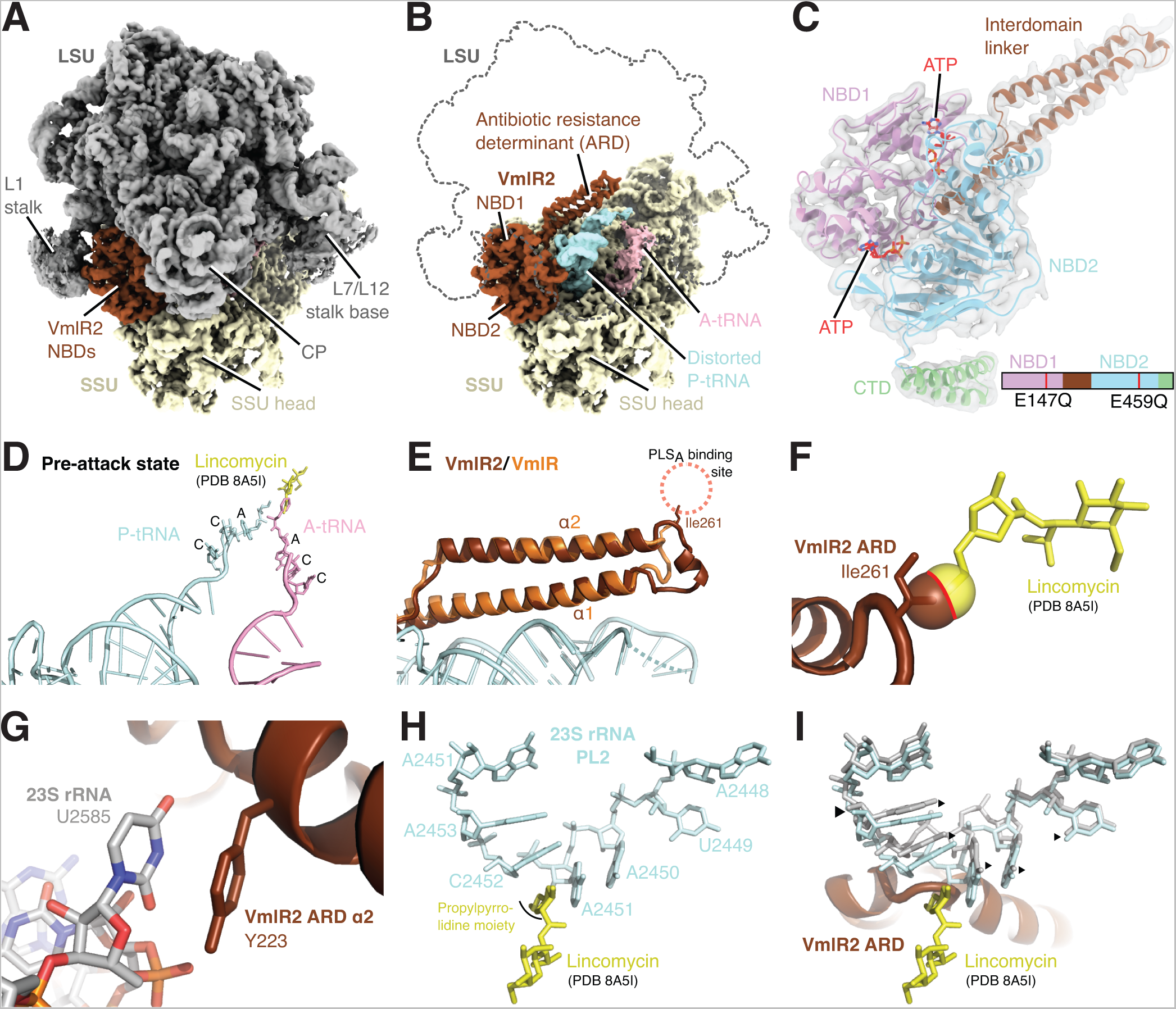
Cryo-EM structure of *N. vireti* VmlR2-EQ_2_ bound to the 70S ribosome. (**A**) Overview of cryo-EM density containing the large ribosomal subunit (grey), VmlR2 (brown), and the small ribosomal subunit (yellow). (**B**) As for **A**, except the large ribosomal subunit is shown as an outline only. A distorted P-tRNA is coloured cyan, and a substoichiometric A-tRNA is pink. (**C**) Isolated density and model of VmlR2 only. The model is coloured by domain, with ATPs coloured red. Non-sharpened maps are shown. (**D**) View of P- and A-tRNAs poised for peptidyl transfer in the PTC (pre-attack state, PDB ID 1VY4, (Polikanov et al., 2014)). The position of lincomycin is superimposed (PDB ID 8A5I, (Koller et al., 2022)). (**E**) Same view as **D** but showing the VmlR2 (brown) and *B. subtilis* VmlR (transparent, orange, PDB ID 6HA8, (Crowe-McAuliffe *et al*., 2018)) interdomain linkers. A dashed circle indicates the site of PLS_A_ binding. Distorted P-tRNAs are shown in cyan, and a dashed line indicates the approximate path of the tRNA CCA-3′ end, which was not modelled. (**F**) Close view of the VmlR2 ARD reaching into the PTC. Ile261 is predicted to clash with superimposed lincomycin (PDB ID 8A5I). Van der Waals radii of the two closest atoms are shown as semi-transparent spheres, with a red line indicating a clash between them. (**G**) VmlR2 Y223 forms a stacking interaction with U2585. (**H**) View of PL2 forming part of the lincomycin binding site with 23S rRNA nucleotides shown in cyan (PDB ID 8A5I). (**I**) Same as **H** but with VmlR2 superimposed. 23S rRNA nucleotides from the VmlR2-bound structure are shown in grey. Models were aligned by 23S rRNA.

As observed previously in ABCF samples prepared by *ex vivo* immunoprecipitation (Crowe-McAuliffe *et al*., 2018; Crowe-McAuliffe *et al*., 2022; Crowe-McAuliffe *et al*., 2021), the density of the P-site tRNA was most consistent with an initiator fMet-tRNA (**Supplementary Figure 5A**), and density for the P-site codon was most consistent with AUG (**Supplementary Figure 5B**). An apparent Shine-Dalgarno–anti-Shine-Dalgarno helix is positioned in the mRNA exit channel, close to, but not visibly contacting, the VmlR2 CTD (**Supplementary Figure 5C**). A sub-stoichiometric tRNA is present in the A site, as observed previously for the 70S-LsaA-EQ_2_ complex from *Enterococcus faecalis,* and sub-classification revealed that the A-site tRNA can adopt multiple conformations (**Figure 2B**, **Supplementary Figure 6**). These observations imply that VmlR2-EQ_2_ had bound to and stalled the 70S ribosome after translation initiation, but before the first translocation event, which is the same state stalled by PLS_A_ and nucleoside antibiotics (Orelle et al., 2013; Polikanov *et al*., 2015).

As expected from the high degree of sequence homology, the structure of VmlR2 bound to the 70S ribosome is globally similar to *B. subtilis* VmlR bound to the 70S, with an root square mean deviation (RMSD) of 1.44 Å between the two factors after least-squares fitting. In particular, the α-helices of the well-resolved interdomain linker, which travel through the path normally occupied by the P-tRNA to reach into the PTC, are highly structurally conserved (**Figure 2D,E**). The ends of the α-helices and the ARD-loop linking them, diverge from the equivalent region in the *B. subtilis* VmlR (**Figure 2E**), as expected from the sequence alignment (**Figure 1B**). Compared to VmlR, the additional eight residues in the VmlR2 ARD are mostly oriented away from the PTC and drug binding site (**Supplementary Figure 7A**), similar to that observed for the ARDs of LsaA, VgaA_LC_, and VgaL, which are also extended compared to *B. subtilis* VmlR despite sharing a similar spectrum of antibiotic resistance (**Supplementary Figure 7B-D**) (Crowe-McAuliffe *et al*., 2021). Indeed, despite containing eight more amino acids than *B. subtilis* VmlR, the VmlR2 ARD reaches a similar site in the PTC as many other PLS_A_-specific ARE-ABCFs (**Figure 2E**, **Supplementary Figure 7**). Residue Ile261 of VmlR2 overlaps with the binding site of PLS_A_ and nucleoside antibiotics, similarly to Phe237 of *B. subtilis* VmlR except inserted approximately 1.4 Å deeper into the PTC (**Figure 2F**, **Supplementary Figure 8A-H**).

We next mutated selected residues of the ARD-loop of VmlR2 to alanine and assessed the ability of the resulting VmlR2 variants to mediate antibiotic resistance in a Δ*vmlR B. subtilis* background (**Table 2**). As observed for other antibiotic-resistance ABCFs, mutating the only residue that overlaps with the PLS_A_ binding site, in this case Ile261 to alanine, did not appreciably shift the antibiotic resistance spectrum of VmlR2 (**Table 1**). However, due to the increased penetration of the VmlR2 ARD into the PTC, an alanine side chain at position 261 is also predicted to still overlap with the PLS_A_ binding site (**Supplementary Figure 8I-K**). We therefore conclude that while the Ile261 sidechain is not required for antibiotic resistance, this experiment is not informative about the role of steric hindrance in the mechanism of antibiotic resistance by VmlR2.

**Table 2.** Effects of CplR and Erm functionality on resistance of C. difficile to nucleoside and PLS_A_ antibiotics. C. difficile MIC testing was carried out in BHIS media, and growth inhibition was scored after 36-48 hours of anaerobic incubation at 37 °C. Plasmid-harboring strains were cultured in BHIS media supplemented with 10 μg/ml thiamphenicol (to maintain the plasmids) and 500 ng/ml anhydrotetracycline (to induce the expression of ARE-ABCF proteins).

|  |  | MIC (µg/ml) |  |  |  |  |  |  |  |  |
| --- | --- | --- | --- | --- | --- | --- | --- | --- | --- | --- |
| *C. difficile* strain | *erm* | A201A | Hygromycin A | Retapamulin | Lincomycin | Clindamycin | Iboxamycin | Virginia mycin M1 | Erythromycin | Florfenicol |
| R20291 | - | 2 | 16 | 0.4 | 16 | 8 | 0.0625 | 128 | >256 | 4 |
| 630 | *c-erm* | 4 | 16 | 0.4 | >256 | 256 | 16 | 128 | >256 | 4 |
| 630Δ*erm* | - | 4 | 16 | 0.8-1.6 | 32 | 16 | 0.25 | 32 | 1 | 4 |
| 630Δ*erm* Δ*cplR* | - | 2 | 16 | 0.1 | 0.5 | 0.125 | 0.0156 | 16 | 1 | 4 |
| 630Δ*erm*/pRFP185 | - | 1 | 8 | 0.2 | 16 | 4 | 0.03125 | 16 | 1 | 4 |
| 630Δ*erm* Δ*cplR*1/pRFP185 | - | 1 | 4 | 0.05 | 0.25 | 0.03125 | 0.03125 | 8 | 1 | 4 |
| 630Δ*erm* Δ*cplR*/pRFP185-*cplR* | - | 2 | 8 | 1.6 | 32 | 8 | <0.00781 | 16 | 1 | 4 |
| 630Δ*erm* Δ*cplR*/pRFP185-*cplR*ΔC | - | 1 | 2 | 0.1 | 1 | 0.5 | 0.0625 | 8 | 1 | 4 |

Mutation of other selected residues close to the drug-binding site resulted in a complex pattern of changes to antibiotic resistance (**Supplementary Figure 9**). Residues Arg259 and Gln260, positioned in the tip of the ARD, were insensitive to mutation to alanine despite making defined contacts with the 23S rRNA (**Supplementary Figure 9A,B**). By contrast, VmlR2 Y266 was required for resistance to tiamulin and virginiamycin M. This residue is positioned similarly to *B. subtilis* VmlR Y240 and VgaA_LC_ Y223, with each residue stacking with 23S rRNA nucleotide U2585 (*E. coli* numbering; **Figure 2G**), and we previously found that a VgaA_LC_ Y223A variant also displayed reduced resistance against PLS_A_ antibiotics (Crowe-McAuliffe *et al*., 2021). The adjacent residue, E265, also had a drastic effect on antibiotic resistance when mutated to alanine. E265 is within hydrogen-bonding distance of 23S rRNA residues A2450 and A2451, which form part of a region we have previously termed PTC loop 2 (PL2, **Supplementary Figure 9C**, (Crowe-McAuliffe *et al*., 2021)). Interactions between VmlR2 and the 23S rRNA in this region may serve to position the VmlR2 ARD in the PTC.

In summary, residues that interact with U2585 and A2450/A2451, located at the beginning of interdomain linker α2, appear to be important for PLS_A_ resistance mediated by VmlR2, while by contrast, mutating the residue that physically overlaps with the PLS_A_ binding site or other residues in the extended tip of the ARD does not affect antibiotic resistance (**Supplementary Figure 9C,D**). These observations are consistent with previous mutagenesis studies of ARE-ABCFs (Crowe-McAuliffe *et al*., 2018; Crowe-McAuliffe *et al*., 2021). Previously, we proposed a model in which ARE-ABCFs confer antibiotic resistance by modulation of the antibiotic binding site (Crowe-McAuliffe *et al*., 2018; Crowe-McAuliffe *et al*., 2022; Crowe-McAuliffe *et al*., 2021). Similar to *B. subtilis* VmlR, VmlR2 modulates the conformation of several 23S rRNA nucleotides that form the PLS_A_ binding site. In particular, nucleotides around A2451, part of PL2, are displaced in the VmlR2-bound structure compared to the antibiotic-bound structures or a stalled elongating ribosome (**Figure 2H,I**, **Supplementary Figure 10**). We therefore propose that binding of VmlR2 to a PLS_A_- or nucleoside-bound ribosome triggers a distortion of the antibiotic-binding site, in turn leading to dissociation of the antibiotic and ultimately allowing translation to resume after dissociation of the ARE-ABCF.

### CplR contributes to intrinsic PLS_A_ resistance in Clostridioides

Next, we characterised the resistance mediated by *C. difficile, C. perfringens* and *C. sporogenes* CplR in the native bacterial hosts.

In our *C. difficile* antibiotic sensitivity experiments we used two strains: the *erm-* R20291 and the *erm+* 630. R20291 is an epidemic hypervirulent strain that belongs to PCR-ribotype 027 (Stabler et al., 2009). The MIC values determined with this strain are directly comparable with *erm-* ATCC 700057 MICs reported by Mitcheltree and colleagues in the original iboxamycin study (Mitcheltree *et al*., 2021). Despite encoding a genomic copy of CplR, both R20291 and ATCC 700057 are sensitive to iboxamycin (MIC of 0.0625 µg/mL for R20291, MIC of 0.25 µg/mL for ATCC 700057 (Mitcheltree *et al*., 2021)) (**Table 2**).

The *C. difficile* 630 wild-type reference strain is a clinical isolate that over the years gained popularity with the research community (Roberts and Smits, 2018). The prophage region of the mobilizable, non-conjugative transposon Tn*5398* present in the genome of *C. difficile* 630 encodes a tandem expression cassette for two identical copies of the *ermB* gene for rRNA adenine N-6-methyltransferase (Farrow *et al*., 2001). This ubiquitous antibiotic resistance determinant confers resistance to macrolides as well as lincomycin and clindamycin (Lai and Weisblum, 1971; Leclercq and Courvalin, 1991). Compared to wild-type 630, the Δ*erm* 630 *C. difficile* is more sensitive to pleuromutilin retapamulin (MIC for of 0.4 µg/mL vs 1.6 µg/mL) as well as lincosamides (MIC for lincomycin of 16 µg/mL vs >256 µg/mL, MIC for clindamycin of 16 µg/ml vs 256 µg/mL and MIC for iboxamycin of 0.25 µg/mL vs 16 µg/mL) (T**able 2**). Since in *B. subtilis* VmlR synergises with Cfr to grant high levels of resistance (Brodiazhenko *et al*., 2022), we next tested the effects of CplR loss in Δ*erm* 630 *C. difficile*. The strain is further sensitised to lincosamides (MIC for lincomycin, clindamycin and iboxamycin of 0.1, 0.5 and <0.125 µg/mL, respectively) as well as virginiamycin M1 (MIC of 16 µg/mL) but not florfenicol or hygromycin A. The effect of CplR expression on nucleoside resistance is minor: a mere 2-fold increase in sensitivity to A201A, and no effect on sensitivity to hygromycin A (**Table 2**, compare 630 Δ*erm* Δ*ARE1* to 630 Δ*erm* as well as 630 Δ*erm* Δ*cplR*/pRFP185 to 630 Δ*erm* Δc*plR*/pRFP185-*cplR*). Our results establish that the synergetic action of Erm and CplR is responsible for the ability of wild-type *C. difficile* 630 to resist PLS_A_ antibiotics, including iboxamycin (MIC of 16 µg/mL).

Next, we tested the effects of CplR loss on antibiotic sensitivity of *C. perfringens* (**Table 3**) and *C. sporogenes* (**Table 4**). Compared to wild-type HN13 *C. perfringens*, the isogenic strain harbouring a markerless disruption of *cplR* is sensitised to the pleuromutilin retapamulin (32-fold increased sensitivity, MIC of 0.8 vs 0.025 µg/mL), lincosamides (MICs dropping 8- (lincomycin), 2- (iboxamycin) or >32-fold (clindamycin)), as well as virginiamycin M1 (a 2-fold MIC drop). Conversely, ectopic overexpression of CplR*^Cp^* from the pJIR 750 vector in the Δ*cplR* background drastically increased the MICs against retapamulin, lincomycin, clindamycin and iboxamycin, exceeding that of the wild-type HN13 *C. perfringens* by 8-fold (**Table 3**).

**Table 3.** Effects of CplR functionality on *C. perfringens* resistance to PLS_A_ and macrolide antibiotics. The MIC test for *C. perfringens* was carried out in GAM media and the growth inhibition was scored after 24 hours of anaerobic incubation at 37 °C. In the case of plasmid-harboring strains GAM media was attentionally supplemented with 10 μg/ml chloramphenicol to maintain the plasmids. Expression of the CplR ARE-ABCF was driven by the native promotor.

| ***C. perfringens* strain** | ***erm*** | **MIC (µg/ml)** |  |  |  |  |  |  |  |
| --- | --- | --- | --- | --- | --- | --- | --- | --- | --- |
|  |  | **A201A** | **Hygromycin A** | **Retapamulin** | **Lincomycin** | **Clindamycin** | **Iboxamycin** | **Virginiamycin M1** | **Erythromycin** |
| wt/pJIR750 | - | 4 | 4 | 0.8 | 4 | 2 | 0.0156 | 3.2 | 1 |
| Δ*cplR*/pJIR750 | - | 4 | 4 | 0.025 | 0.25 | <0.0625 | 0.0078 | 1.6 | 1 |
| Δ*cplR*/pJIR750-*cplR* | - | 8 | 8 | 12.8 | 32 | 16 | 0.125 | 3.2 | 1 |

**Table 4.** Effects of CpiR functionality on C. sporogenes resistance to nucleoside, PLSA and macrolide antibiotics. The MIG test for C. sporogenes was carried out in GAM media, and the growth inhibition was scored after 24 hours of anaerobic incubation at 37 °C.

| *C. sporogenes*<br>strain | *erm* | MIC (µg/ml) |  |  |  |  |  |  |  |
| --- | --- | --- | --- | --- | --- | --- | --- | --- | --- |
|  |  | A201A | Hygromycin A | Retapamulin | Lincomycin | Clindamycin | Iboxamycin | Chloramphenicol | Erythromycin |
| JCM1416 wt | - | 1 | 4 | 25.6 | 32 | 16 | 0.125 | 2 | 0.5 |
| *cplR::intron-ermB* | *c-ermB* | 1 | 2 | 0.025 | 256 | 8 | 0.031<br>25 | 2 | >256 |
| *sigD::intron-ermB* | *c-ermB* | 1 | 4 | 25.6 | 256 | 128 | 16 | 2 | >256 |

To test the role of CplR in antibiotic resistance of *C. sporogenes*, we constructed a strain in which the *cplR^Cs^* gene is disrupted by the insertion of the *intron*-*ermB* resistance marker (**Table 4**). To deconvolute the effect of the introduced *ermB* gene on antibiotic resistance from that of *cplR*^Cs^ disruption, we also constructed a strain in which the *intron*-*ermB* marker is introduced into the *sigD* locus. Compared to the isogenic wild type, the *cplR*::*intron-ermB* strain is more sensitive to retapamulin (MICs of 0.025 µg/mL vs 25.6 µg/mL, **Table 4**) suggesting that CplR mediates resistance to pleuromutilins. As expected, to the introduction of the ErmB marker the *cplR*::*intron-ermB* strain displays higher resistance to lincomycin than the isogenic wild type (MIC for lincomycin of 256 µg/mL vs 32 µg/mL, **Table 4**). At the same strain, the *cplR*::*intron-ermB* strain is more sensitive to iboxamycin (MICs of 0.031 µg/mL vs 0.125 µg/mL), consistent with the role of CplR in lincosamide resistance. The antibiotic resistance function of ClpR was is further supported by the clindamycin and iboxamycin MIC of the *cplR*-disrupted mutant being lower than that of the *sigD* mutant (compare *cplR*:*intron-ermB* to *sigD*::*intron-ermB*: clindamycin MICs of 8 µg/mL vs 128 µg/mL, iboxamycin MICs of 0.031 µg/mL vs 16 µg/mL) (**Table 4**). Finally, the effects on nucleoside antibiotic sensitivity are minor (2-fold increased sensitivity to hygromycin B, and no effect on sensitivity to A201A).

Taken together, our results establish ARE1 CplR as a Clostridiales genome-encoded PLS_A_ resistance determinant and suggest that high PLS_A_ resistance levels of *C. difficile* 630 are mediated through a combined action of direct ribosome protection by CplR and 23S rRNA modification by the ErmB methyltransferase.

### An antibiotic challenge induces the accumulation of *C. difficile* CplR full-length mRNA, and expression is re-repressed once bacterial growth is restored

Expression of ARE-ABCFs is typically induced upon a sub-MIC antibiotic challenge through a de-repression mechanism, with the full-length mRNA levels being kept low in the absence of the cognate antibiotic due to premature transcription termination in the absence of ribosomal stalling on a regulatory uORF (Cai *et al*., 2021; Dar *et al*., 2016; Koberska *et al*., 2021; Ohki *et al*., 2005; Takada *et al*., 2022; Vimberg *et al*., 2020).

Genome-wide mapping of transcriptional start sites (TSSs; identified using dRNA-seq) and termination sites (TTSs; identified using RNAtag-seq) in *C. difficile* 630 (Fuchs *et al*., 2021) revealed that i) the *cplR* ORF preceded by a 212 nucleotide-long 5ʹ upstream region (CDIF630nc_084), and that ii) as expected for an ARE-ABCF, under non-inducing conditions, the transcript prematurely terminates at start codon of the *cplR* ORF (CDIF630_02847) (**Figure 3A**). Since antibiotic-induced ribosomal stalling by a regulatory uORF is a common regulatory strategy employed for inducible control of ARE-ABCF expression (Cai *et al*., 2021; Dar *et al*., 2016; Koberska *et al*., 2021; Ohki *et al*., 2005; Takada *et al*., 2022; Vimberg *et al*., 2020), we have mounted an *in silico* search for regulatory ORFs of CplR and close relatives using our bioinformatic tool for uORF detection, uORF4u (Egorov and Atkinson, 2022). Sequence conservation analysis reveals that the uORF of Clostridial CplR has a consensus of MR[I/M], with *C. difficile cplR* uORF specifically encoding Met-Arg-Ile (**Figure 3B,C**). The consensus is different from the uORF4u-computed uORF for VmlR (MIN, **Figure 3C**), VmlR2 (MK[L/Q], **Figure 3D**) and LsaA (MAGN, **Figure 3F**), despite the four ARE-ABCFs having similar resistance spectra. The identified short uORFs are likely to be stalled by PLS_A_ and nucleoside antibiotics, which inhibit translation after initiation with little context-specificity (Orelle *et al*., 2013; Polikanov *et al*., 2015). We could not identify an adequate number of sufficiently closely related homologues for the tool to be able to predict any conserved uORFs for Ard1.

**Figure 3.**
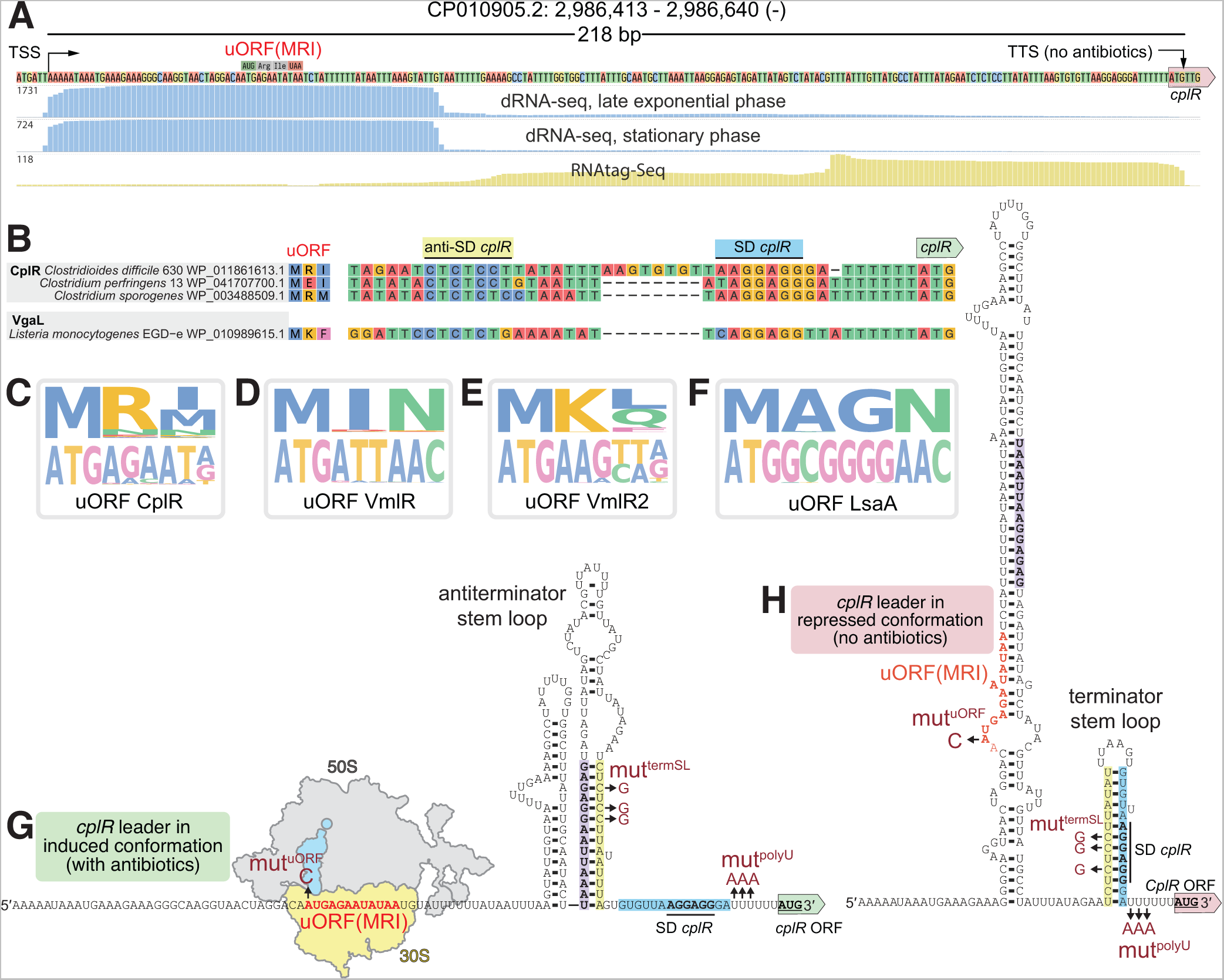
Nucleotide sequence and predicted secondary structures of the 5′ leader region of *CplR* mRNA in induced and repressed conformations. (**A**) dRNA-seq (sampled in late exponential or stationary phase) and RNAtag-seq reads for annotation of transcriptional start sites (TSSs) and termination sites (TTSs), respectively, mapped to 5ʹ upstream region (CDIF630nc_084) preceding the *cplR* ORF (CDIF630_02847) (Fuchs *et al*., 2021). (**B**) Alignments of 5ʹ upstream regions elements in Clostridial CplRs and *L. monocytogenes* Lmo0919/VgaL focused on the uORF and anti-Shine-Dalgarno (anti-SD) elements. (**C**-**F**) Consensus sequences for (**C**) CplR, (**D**) VmlR, (**E**) VmlR2 and (**F**) LsaA uORF elements. The full uORF peptide alignments are shown in **Supplementary Table 3**. (G and H) Predicted secondary structure of *cp/R* mRNA 5’ leader region in repressed (G, in the absence of antibiotic-induced ribosomal stalling on the uORF(MRI)) and induced (H, in the presence of ribosomal stalling). Substitutions that were used for experimental probing the key functional elements are shown in red.

We predicted secondary structures of the 5ʹ *cplR* leader region in induced (i.e. full-length mRNA with 70S ribosome stalled on the uORF in the presence of antibiotics) and repressed (i.e. prematurely terminated mRNA generated in the absence of antibiotics) conformations using RNAfold (Gruber *et al*., 2008) and Mfold (Zuker, 2003). In the repressed conformation the 5ʹ *cplR* leader region is predicted to form two stable stem loop structures (**Figure 3G**). The 5ʹ hairpin sequesters the putative regulatory uORF(MRI). The 3ʹ terminator hairpin is followed by a single-stranded polyU stretch, thus constituting a canonical signal for efficient intrinsic termination in bacteria (Farnham and Platt, 1981; Gusarov and Nudler, 1999; Sipos et al., 2007; Yarnell and Roberts, 1999). Importantly, the stem loop of the terminator hairpin sequesters the Shine-Dalgarno element positioned to drive the expression of the *cplR* ORF, potentially mediating the suppression of the *cplR* expression from full-length mRNA transcripts that fail to terminate prematurely. In repressed conformation the terminator hairpin is not formed, with an anti-terminator stem loop formed instead which allows transcription of full-length mRNA (**Figure 3H**). Furthermore, the Shine-Dalgarno that drives the *cplR* expression is readily accessible for initiating ribosomes (**Figure 3D**).

To test antibiotic-inducible expression of *C. difficile cplR,* we cultured 630Δ*ermR C. difficile* in the presence of increasing sub-MIC concentrations of retapamulin (2-200 ng/mL; MIC of 800-1600 ng/mL) and lincomycin (0.02-2.0 µg/mL; MIC of 32 µg/mL) and assessed the levels of *cplR* mRNA by RT-PCR using two primer pairs: one amplifying 5ʹ leader region and the other targeting the ORF (**Figure 4A**). As expected, the level of *cplR* mRNA was increased in a dose-dependent manner for both retapamulin and lincomycin treatment, with the RT-PCR signal for both mRNA regions increasing upon antibiotic challenge (**Figure 4B**).

**Figure 4.**
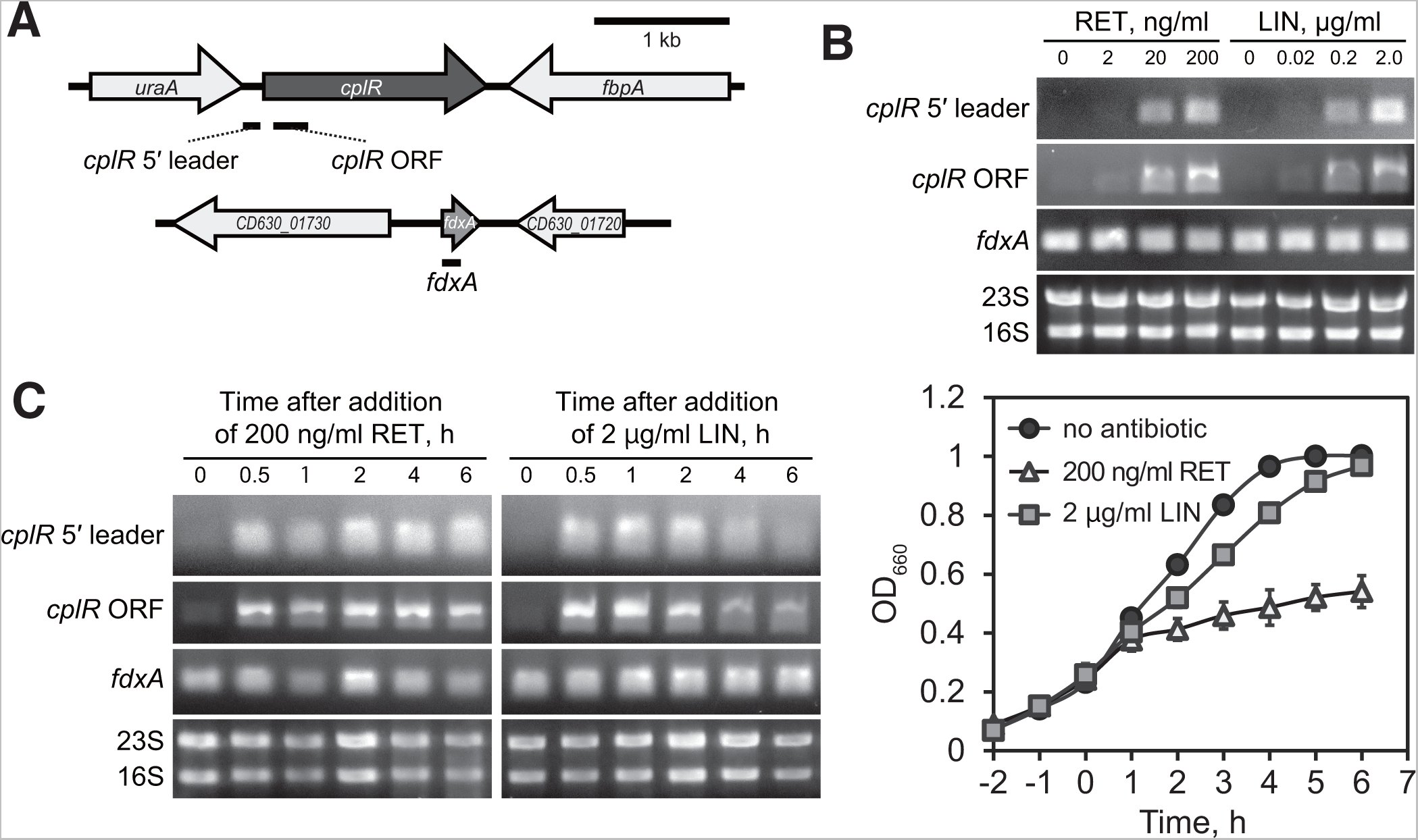
*cplR* expression is de-repressed in *C. difficile* upon retapamulin or lincomycin challenge and re-repressed once the bacterial growth recovers. (**A**) Experimental setup used for *cplR* mRNA detection by RT-PCR using primers specific for *cplR* 5′ leader region, *cplR* ORF and *fdxA* ORF. (**B**) *cplR* mRNA accumulates in a dose-dependent manner in response to the addition of increasing concentrations of either retapamulin (RET) or lincomycin (LIN). The antibiotics were added to logarithmic phase *C. difficile* 630 Δ*ermR* cultures of and total RNA collected 2 hours after the antibiotic challenge. 23S and 16S rRNA were used as loading controls. (**C**) Kinetics of *cplR* mRNA accumulation in response to a sub-MIC challenge with retapamulin (200 ng/ml) or lincomycin (2 µg/ml). (*Left*) *cplR* 5′ leader region, *cplR* ORF and *fdxA* ORF were probed by RT-PCR amplification. (*Right*) Growth kinetics of *C. difficile* in the absence or presence of sub-MIC concentrations of retapamulin (200 ng/ml) or lincomycin (2 µg/ml). The time point of antibiotic addition is set as zero on the time axis.

Next, we investigated the time-course of *cplR* expression upon antibiotic challenge. We hypothesised that when the inhibitory effect of the antibiotic on translation is relieved by the ARE-ABCF, the expression of the factor should subside as the antibiotic-induced ribosomal stalling that drives the expression is abolished. To test this hypothesis, we contrasted the RT-PCR data with bacterial growth kinetics. The *cplR* expression quickly responded to the antibiotic challenge, peaking within 30 minutes after the addition of antibiotics (**Figure 4C**, *left*). Importantly, while the retapamulin-induced growth inhibition was severe and persisted for the whole duration of the experiment, the lincomycin-induced growth inhibition is much milder and at the later time points the growth kinetics are similar to that one the untreated control (**Figure 4C**, *right*). Consistent with the prediction, while in the presence of 200 ng/mL retapamulin the maximal levels of *cplR* mRNA are maintained throughout the 6 hours of the RT-PCR time course, but the *cplR* mRNA levels gradually decreased as bacterial growth – and, by inference, protein synthesis – has recovered from the lincomycin challenge (**Figure 4C**, *left*).

Collectively, our results suggest that i) inhibition of translation by retapamulin or lincomycin induces accumulation of *cplR* mRNA and ii) induction of *cplR* mRNA is abrogated once expression of the factor overcomes the drug-induced translation inhibition.

### The 5′ leader region of *cplR* controls the induction of ABCF expression upon an antibiotic challenge

To test the role of the predicted 5ʹ *cplR* leader region elements in antibiotic-induced expression of *cplR* in *C. difficile* we used a series of reporter constructs in which the expression of the mCherry red fluorescent protein was driven by the native *cplR* promoter (P*_cplR_*) with either wild-type or mutated 5ʹ leader variants (**Figure 5A**). We used i) RT-PCR to assess the effects of 5ʹ leader mutations and/or antibiotic challenge that act on the level of transcription and ii) mCherry fluorescence measurements to assess the compound effects that act both on transcriptional and translational levels simultaneously.

**Figure 5.**
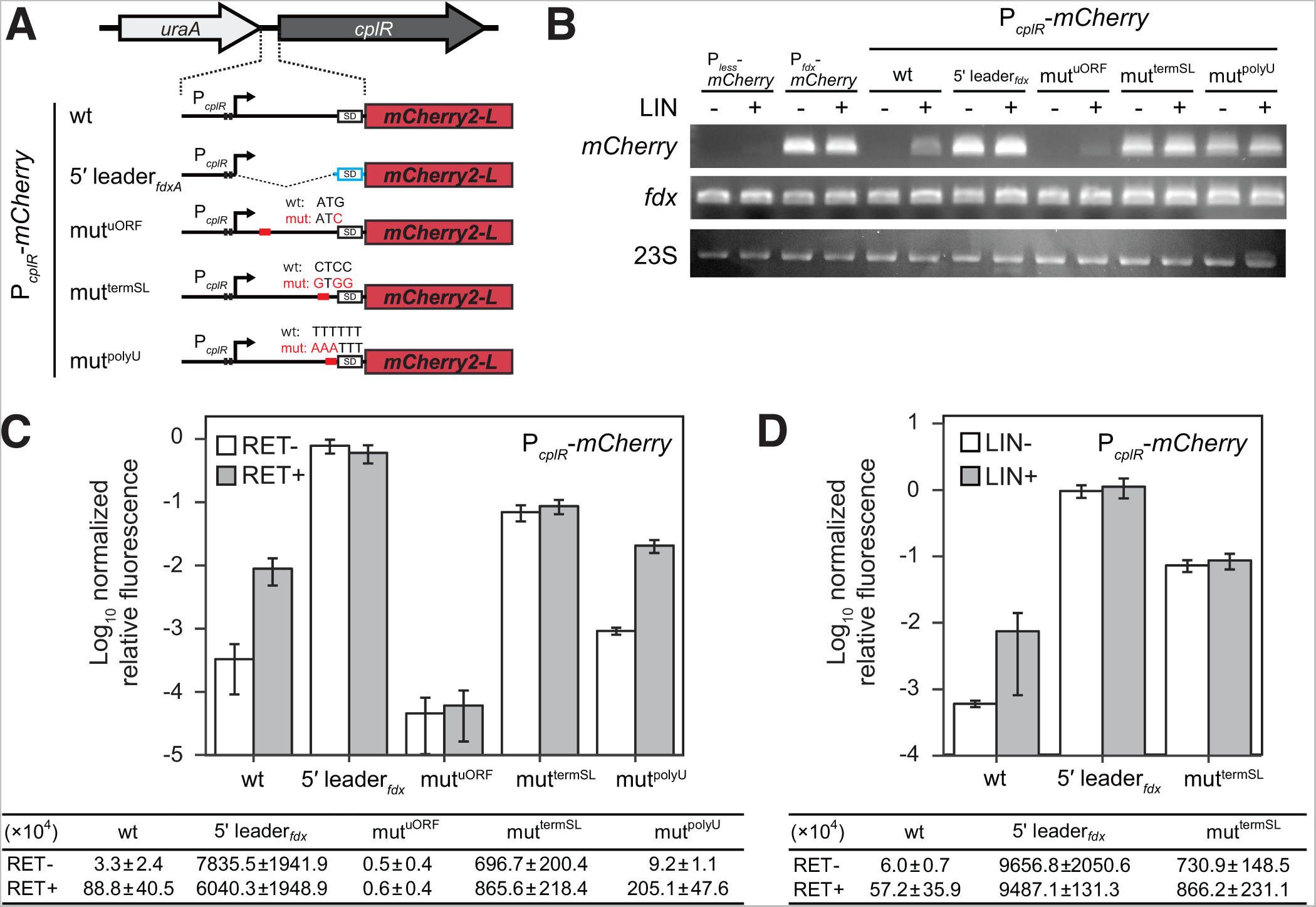
The ribosomal stalling on the uORF but not the premature RNAP termination is essential for induction of *ARE1* in response to antibiotic challenge. (**A**) Experimental setup used for mutational probing of the *cplR* 5′ leader region. Fluorescent reporters relied on mCherry expression driven by the *cplR* promotor (P*_CplR_*) using wither i) wild-type *cplR* 5′ leader, ii) *fdxA* 5′ leader (5′ leader*­*), iii-v) *cplR* 5′ leader with substitutions in the uORF start codon (mut^uORF^), the intrinsic terminator stem (mut^termSL^) or the polyU track downstream of the intrinsic terminator stem loop (mut^polyU^). Locations of the substitutions on the mRNA secondary structure are presented on **Figure 3CD**. (**B**) Accumulation *mCherry* mRNA variants with different *cplR* 5′ leaders in response to a sub-MIC challenge with lincomycin (2 µg/ml). The antibiotics were added to the logarithmic phase of *C. difficile* 630 Δ*ermR* culture and total RNA was collected 30 mins after the antibiotic challenge. *mCherry* ORF and *fdx* ORF were probed by RT-PCR amplification. 23S rRNA was used as loading control. (**C** and **D**) Mutational probing of the *cplR* 5′ leader. Expression of mCherry driven by is 5′ leader*_fdxA_* (P*_fdx_*-*mCherry*) is insensitive to sub-MIC challenge with retapamulin (**C**) or lincomycin (**D**). Wild-type P*_cplR_*-*mCherry* reporter is induced by both retapamulin (**C**) and lincomycin (**D**), though the signal is much lower than in the case of P*_fdx_*-*mCherry*. The mut^termSL^ reporter is constitutively induced and is insensitive to addition of either retapamulin (**C**) or lincomycin (**D**). The mut^polyU^ reporter is repressed in the absence of antibiotics and is induced by the retapamulin challenge (**C**). To calculate the relative promoter activity, relative fluorescence intensity normalised by O.D._600_ was further normalised by the signal from the constitutive P*_fdx_-mCherry* reporter.

When P*_cplR_* is used to drive the expression of *mCherry* preceded by wild-type *cplR* 5ʹ leader, while virtually no RT-PCR or fluorescence signal is detectable in the absence of antibiotics, whereas addition of either retapamulin (at 0.2 µg/µL) or lincomycin (at 2 µg/mL) results in strong induction of the reporter (**Figure 5BC**, **Supplementary Figure 11**). Conversely, when the 5ʹ leader region is replaced with that of constitutively and strongly expressed ferredoxin gene (*fdx*) (Takamizawa et al., 2004), both the RT-PCR (**Figure 5B**) and fluorescence (**Figure 5C,D**) signal drastically increased and became insensitive to the antibiotic challenge. Taken together, these results suggest that our mCherry-based reporter strategy faithfully replicates the regulatory mechanisms that natively control the expression of *cplR*.

After establishing the validity of the assay, we next introduced point mutations in the *cplR* 5ʹ leader probing i) the regulatory uORF (AUG to AUC substitution of the initiation codon; mut^uORF^), ii) the terminator stem loop that sequesters the *cplR* main ORF Shine-Dalgarno motif (triple C-to-G substitution disrupting the helix; mut^termSL^) and, finally, iii) the terminator polyU tract (triple U-to-A substitution; mut^polyU^) (**Figures 3G,H 5A**).

The mut^uORF^ fluorescent reporter is inactive both in the presence and the absence of either retapamulin (**Figure 5C**) or lincomycin (**Figure 5D**), which is consistent with translation of the uORF being essential for the induction of *cplR.* The mut^termSL^ reporter in which premature transcription termination is ablated and the Shine-Dalgarno (SD) element that drives expression of *cplR* is made directly accessible for translation initiation is i) constitutively active and non-responsive to addition of antibiotics, and ii) the fluorescent signal is more than 10-fold higher than that of the fully induced wild-type 5ʹ leader reporter construct (**Figure 5CD**). By contrast, the expression of mCherry from the mut^polyU^ reporter is i) tightly regulated by antibiotics, with only very weak signal detectable in the absence of retapamulin, ii) in the presence of retapamulin, the fluorescent signal was about twice (2.3-fold) as strong as that in the case of the wild-type reporter (**Figure 5C**). In the case of the mut^polyU^ reporter the full-length mRNA is expected to be constitutively produced (just as in the case of mut^termSL^). Our RT-PCR experiments directly support this prediction: in good agreement with consistent production of the mut^polyU^ reporter mRNA, the maximum induced signal is higher for the mut^polyU^ reporter than for the wild-type reporter (**Figure 5B**). At the same time, the *cplR* SD element is expected – just as in the wild-type reporter – to be accessible only upon ribosomal stalling on the uORF, and, evidently, translational attenuation is sufficient to ensure the suppression of *cplR* expression in the absence of antibiotic.

Collectively, our results establish that translational attenuation and transcriptional attenuation synergise to control the inducible expression of CplR in Clostridiales.

## DISCUSSION

We have experimentally characterised five genome-encoded ARE-ABCF proteins that are implicated in the intrinsic antibiotic resistance of *S. capreolus* (AAF1 Ard1), *N. vireti* (ARE2 VmlR2), and Clostridioides *C. difficile, C. perfringens* and *C. sporogenes* (ARE1 CplR). As we show here, the Δ*vmlR B. subtilis* strain is a convenient surrogate host for uncovering the resistance spectrum of different ARE-ABCFs. Thus, antibiotic sensitivity assays using heterologous expression in Δ*vmlR B. subtilis* could be used to study other chromosomal ABCFs, guided by bioinformatic surveys.

Our cryo-EM reconstruction of the *N. vireti* VmlR2-70S complex is broadly similar to structures of other ARE-ABCFs with specificity for PLS_A_ antibiotics. The ARD of these elements is counter-intuitively both essential for resistance and poorly conserved at the sequence level, exemplified by the eight-amino-acid insertion in the VmlR2 ARD compared to the *B. subtilis* VmlR ARD, despite the similar antibiotic resistance profiles of both these ARE-ABCFs (**Figure 1B**, **Table 1**, **Supplementary Figure 7**). We observe the extended VmlR2 ARD to be oriented mostly away from the PTC and towards the P-tRNA, reminiscent of the ARDs from more distantly-related PLS_A_-specific ARE-ABCFs such as LsaA, VgaA_LC_, and VgaL (**Supplementary Figure 7**). Thus, ARDs from PLS_A_-specific ARE-ABCFs can accommodate addition or deletion of amino acids while only modestly modulating their position with respect to the PTC. To date, all ribosome-bound ARE-ABCF structures have shown either a modest overlap between the factor and the antibiotic binding site, or no overlap at all—a pattern which holds for ARE-ABCFs specific for both PLS_A_s as well as macrolide, ketolide, and streptogramin Bs (Crowe-McAuliffe *et al*., 2018; Crowe-McAuliffe *et al*., 2021; Mohamad *et al*., 2022; Su *et al*., 2018). This pattern is challenged by our results expanding the specificity of VmlR and VmlR2 to the nucleoside antibiotics A201A and Hygromycin A (**Table 1**), as each of these antibiotics occupy the PLS_A_ binding site but additionally extend towards the A site. Modelled overlaps between VmlR and these antibiotics are extensive (**Supplementary Figure 8**).

The most clinically important aspect of this work is the detailed study of the Clostridial ARE1 CplR. The Gram-positive anaerobic spore-forming bacterium *C. difficile* is a causative agent of nosocomial and chronic infections as well as healthcare-associated diarrhoea (Czepiel et al., 2019). Disruption of the protective gut microbiota by broad-spectrum antibiotics allows for efficient proliferation of antibiotic-resistant *C. difficile* in the gut, thus triggering *C. difficile* infection, CDI (Martin et al., 2016). *C. difficile* has high levels of antibiotic resistance, and treatment with lincosamides such as clindamycin has long been recognised as a risk factor for development of CDI, especially in the case of *erm*+ strains (Bishara *et al*., 2008; Buffie *et al*., 2012; Duffy *et al*., 2020; Johnson *et al*., 1999; Sullivan *et al*., 2001). As we show here, i) similarly to *L. monocytogenes* ARE-ABCF VgaL and 23S ribosomal RNA methyltransferase Cfr (Brodiazhenko *et al*., 2022), *C. difficile* CplR synergises with the 23S rRNA methyltransferase Erm to grant high levels of PLS_A_ resistance, and, ii) analogously to other experimentally studied ARE-ABCFs, induction of CplR is controlled by ribosomal stalling on a regulatory uORF. These results warrant a systematic exploration for co-occurrence of ARE-ABCFs and 23S rRNA methyltransferase antibiotic resistance determinants, with a special focus on the cases of their co-localisation in one operon on a mobile element (as is observed for LsaA and Cfr (Gumkowski *et al*., 2019; Kehrenberg *et al*., 2007)). We demonstrate the utility of our bioinformatic tool uORF4u (Egorov and Atkinson, 2022) for identification of regulatory uORFs and their consensus sequences. The induction of ARE-ABCFs relies on antibiotic-induced ribosomal stalling on regulatory uORFs (Cai *et al*., 2021; Dar *et al*., 2016; Koberska *et al*., 2021; Ohki *et al*., 2005; Takada *et al*., 2022; Vimberg *et al*., 2020), and for elongation inhibitors such as macrolides, phenicols and oxazolidinones, antibiotic-induced ribosomal stalling is context-specific (Beckert *et al*., 2021; Davis et al., 2014; Kannan et al., 2014; Seip et al., 2018; Syroegin et al., 2022a; Syroegin *et al*., 2022b; Tsai *et al*., 2022). A dedicated follow-up investigation of the relationship between uORF consensuses and ARE-ABCF resistance and antibiotic inducibility spectra is warranted. The specific sequence conservation of the uORF could point towards as-yet-uncharacterized patterns of sequence-specific antibiotic-mediated ribosomal stalling that are used by the cell to trigger the resistance factor induction upon an antibiotic challenge. As we have shown earlier, in the case of *B. subtilis* VmlR, the sequence identity of the regulatory uORF(MIN) is crucial for efficient induction upon a lincomycin challenge (Takada *et al*., 2022), despite this antibiotic not being known to act in a context-dependent manner. These insights, in turn, could potentially be leveraged for development of antibiotic variants with unnatural novel stalling profiles that are not detected by uORF-mediated attenuation mechanisms, and, therefore, do not trigger inducible resistance.

## SUPPLEMENTARY DATA

**Supplementary Data** are available at bioRxiv.

## ACKNOWLEDGMENTS

We thank Michael Hall for help with cryo-EM data collection and grid preparation, Farren Isaacs for providing the pRK24 plasmid (Ma *et al*., 2014) as well as Robert Fagan and Neil Fairweather for providing the pRPF185 plasmid. The electron microscopy data was collected at the Umeå Core Facility for Electron Microscopy, a node of the Cryo-EM Swedish National Facility, funded by the Knut and Alice Wallenberg, Family Erling Persson and Kempe Foundations, SciLifeLab, Stockholm University and Umeå University.

---

**Supplementary Table 1. Strains, plasmids and oligonucleotides used in this study.**

The table is provided as a separate excel file. Details of plasmid and stain construction are also provided in the **Supplementary Table 1.** *B. subtilis* strains used in the study were trpC2 *!wm/R* (Crowe-McAuliffe *eta/.,* 2018).

**Supplementary Table 2.**
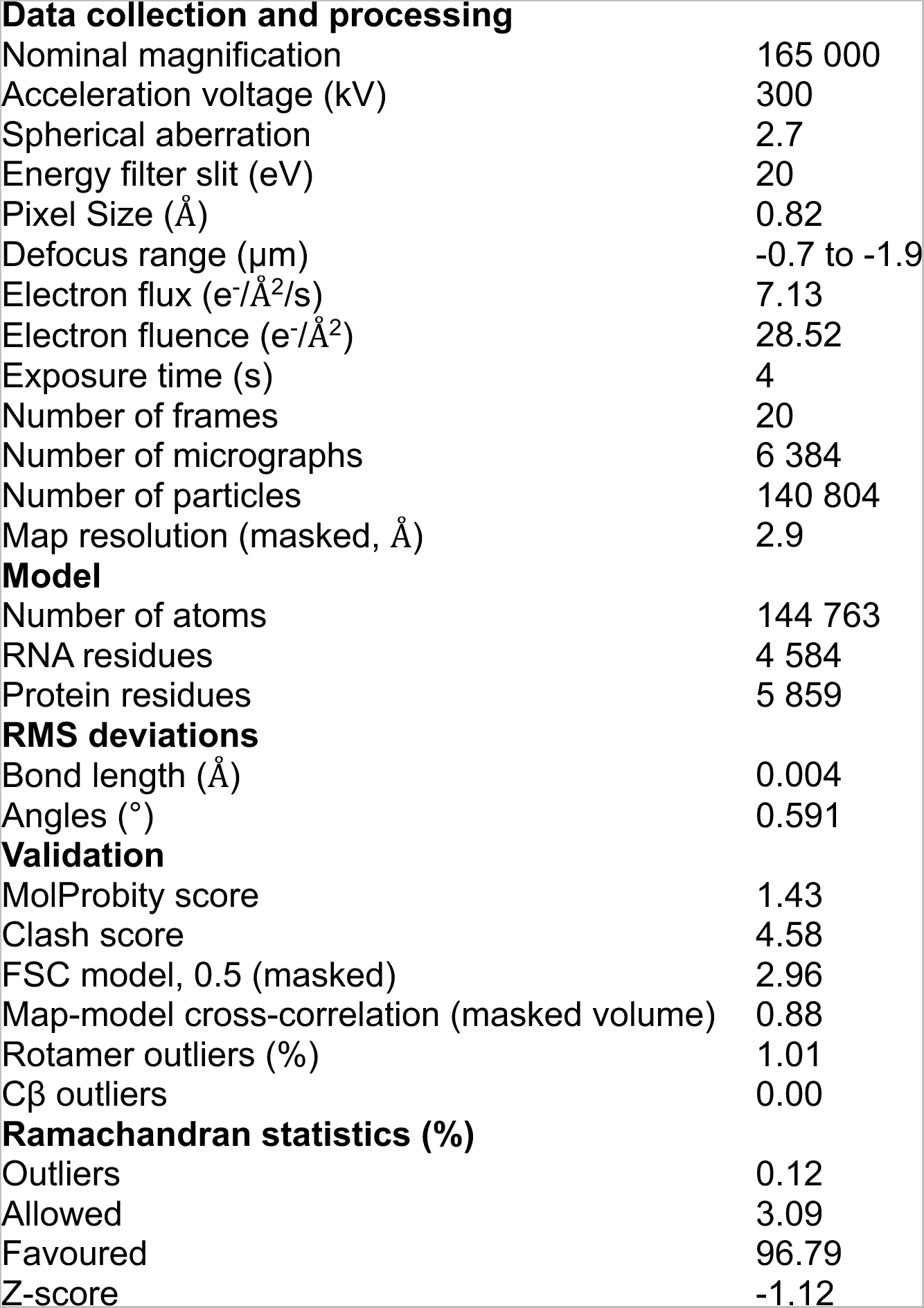
Cryo-EM data collection, modelling and refinement statistics.

|  |  |
| --- | --- |
| Nominal magnification | 165 000 |
| Acceleration voltage (kV) | 300 |
| Spherical aberration | 2.7 |
| Energy filter slit (eV) | 20 |
| Pixel Size (Å) | 0.82 |
| Defocus range (µm) | -0.7 to -1.9 |
| Electron flux (e<sup>-</sup>/Å<sup>2</sup>/s) | 7.13 |
| Electron fluence (e<sup>-</sup>/Å<sup>2</sup>) | 28.52 |
| Exposure time (s) | 4 |
| Number of frames | 20 |
| Number of micrographs | 6 384 |
| Number of particles | 140 804 |
| Map resolution (masked, Å) | 2.9 |
| **Model** |  |
| Number of atoms | 144 763 |
| RNA residues | 4 584 |
| Protein residues | 5 859 |
| **RMS deviations** |  |
| Bond length (Å) | 0.004 |
| Angles (°) | 0.591 |
| **Validation** |  |
| MolProbity score | 1.43 |
| Clash score | 4.58 |
| FSC model, 0.5 (masked) | 2.96 |
| Map-model cross-correlation (masked volume) | 0.88 |
| Rotamer outliers (%) | 1.01 |
| Cβ outliers | 0.00 |
| Outliers | 0.12 |
| Allowed | 3.09 |
| Favoured | 96.79 |
| Z-score | -1.12 |

**Supplementary Table 3. Multiple sequence alignments of ARE-ABCF leader peptides.**

The table is provided as a separate excel file. Complete multiple sequence alignments of leader peptides that regulate expression of CpiR, VmiR, VmiR2, and LsaA generated by uORF4u (Egorov and Atkinson, 2022).

**Supplementary Figure 1.**
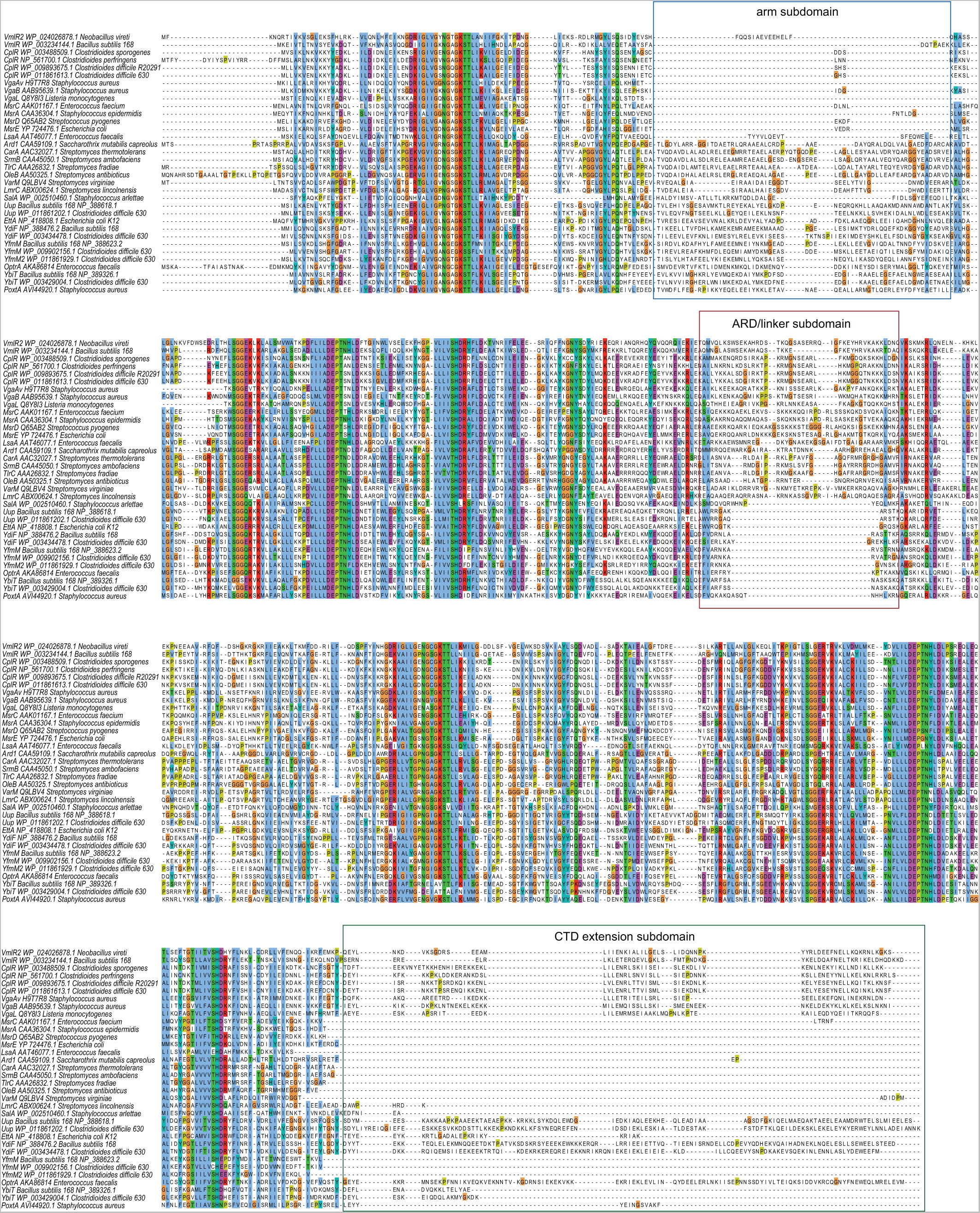
Multiple sequence alignment of selected ABCF proteins. Boxes show the three variable subdomains: the arm, ARD (also referred to as linker), which separates the two conserved ABC domains, and C-terminal domain (CTD).

**Supplementary Figure 2.**
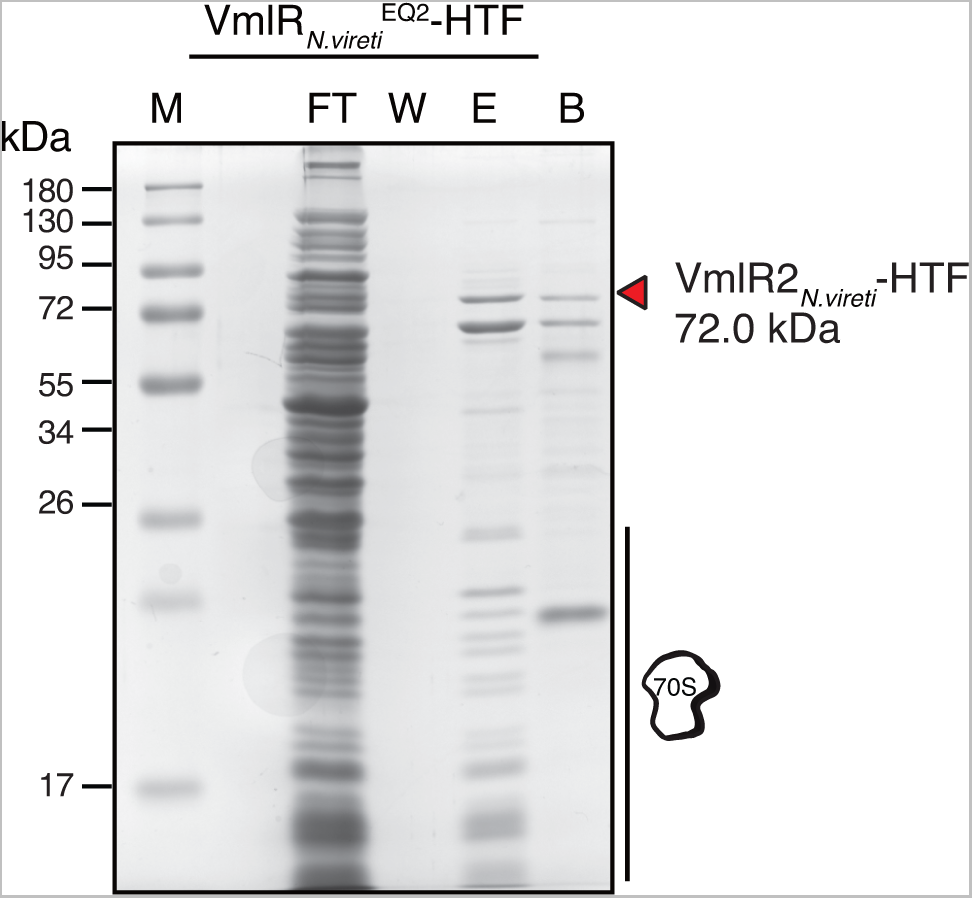
Preparation of samples for cryo-EM reconstructions. Affinity purification of C-terminally HTF-tagged VmlR2*_N.vireti_*^EQ2^ expressed in *B. subtilis* VHB220 (Δ*vmlR thrC*::P*_hy-spnak_* -vmlR2*_N.vireti_*^EQ2^*-HTF*; HTF stands for His_6_-TEV-FLAG_3_). Samples: marker: 2 μL of molecular weight marker; flowthrough: 10 μL; wash: 10 μL of last wash before specific elution; elution: 10 μL of elution with FLAG_3_; beads: 2 μL of 2xSDS loading buffer boiled with 50 μL of beads. The samples were resolved on 12% SDS-PAGE gel.

**Supplementary Figure 3.**
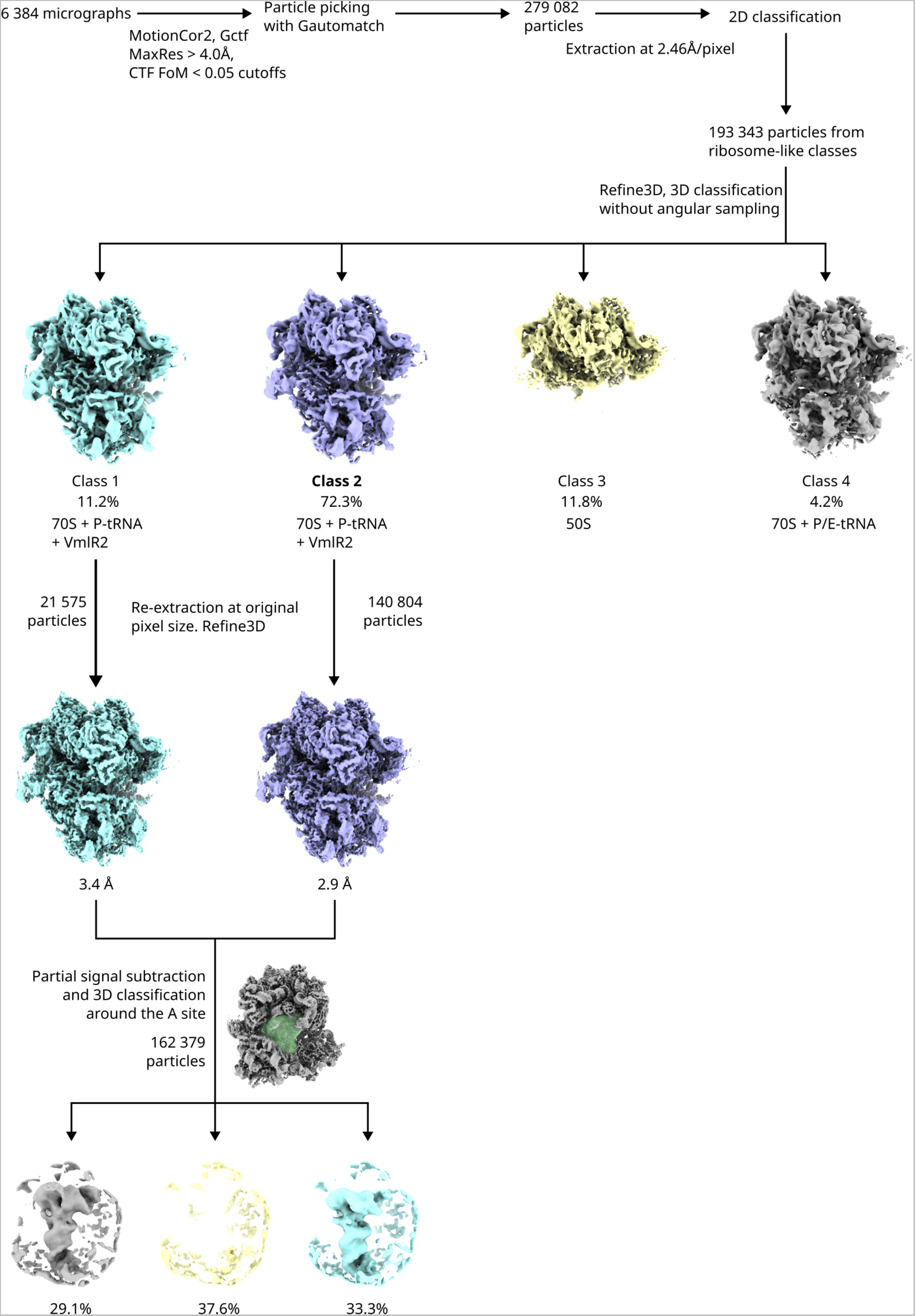
Cryo-EM processing scheme for VmiR2-705. See also methods for details.

**Supplementary Figure 4.**
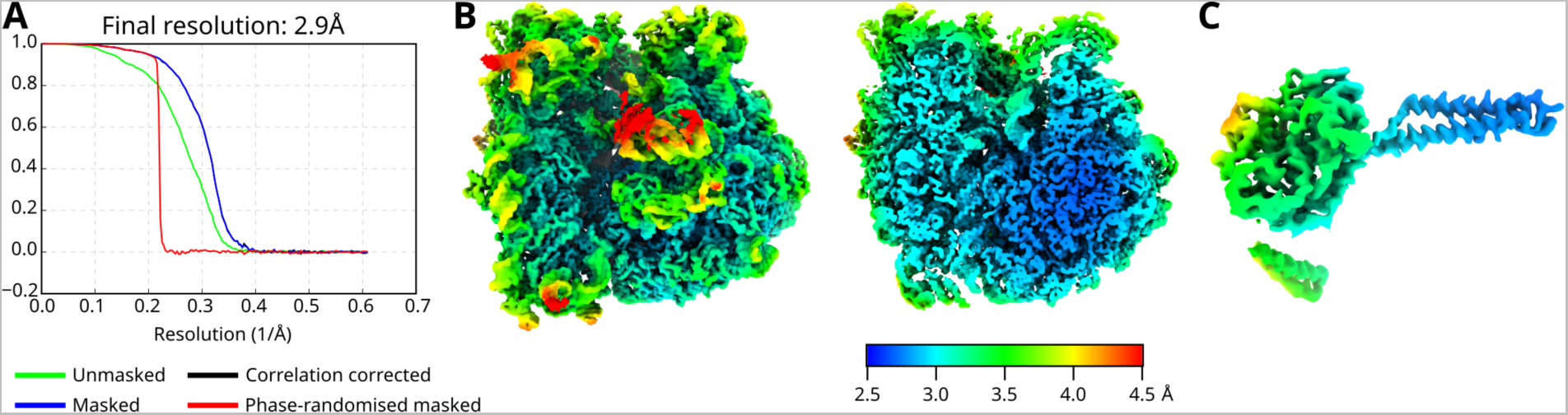
FSC curves and local resolution of the VmlR2-EQ_2_-70S volume. (**A**) FSC curves. (**B**) Whole volume (left) and cut-through (right) coloured according to local resolution. (**C**) Isolated density for VmlR2 coloured according to local resolution. Non-sharpened maps are shown.

**Supplementary Figure 5.**
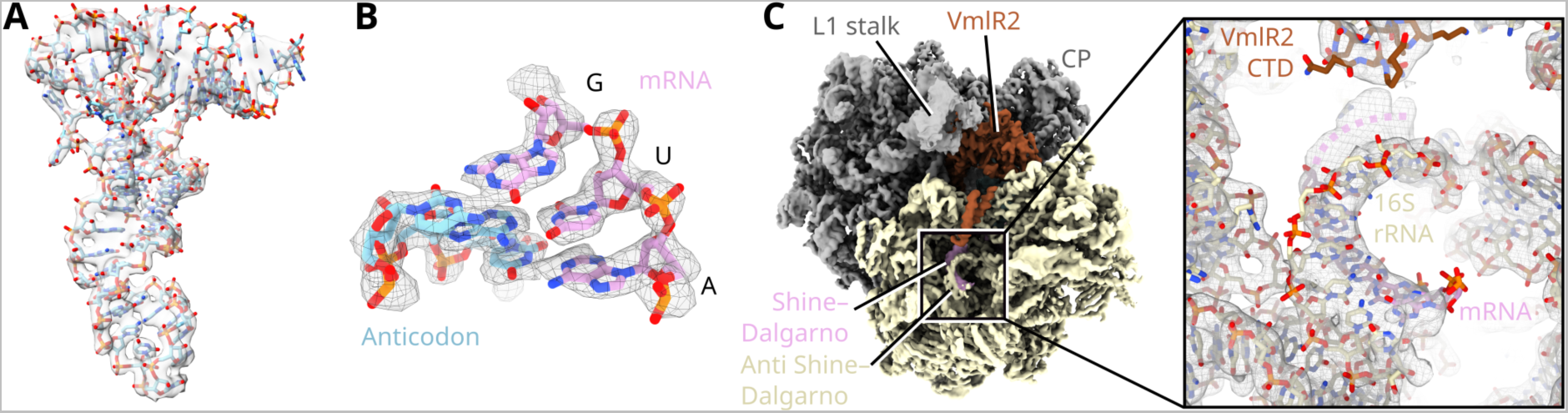
VmlR2 binds an initiation complex. (**A**) Isolated density and model of the distorted P-tRNA. The modelled residues correspond to an fMet-initiator tRNA. The CCA-3′ end is poorly resolved and not included in the model. (**B**) Isolated density and model for the codon-anticodon interaction. The codon is modelled as AUG (pink). The anticodon is shown in blue. (**C**) View of the VmlR2-70S complex focusing on the mRNA exit channel. Density corresponding to a putative Shine-Dalgarno—anti-Shine-Dalgarno interaction is indicated.

**Supplementary Figure 6.**
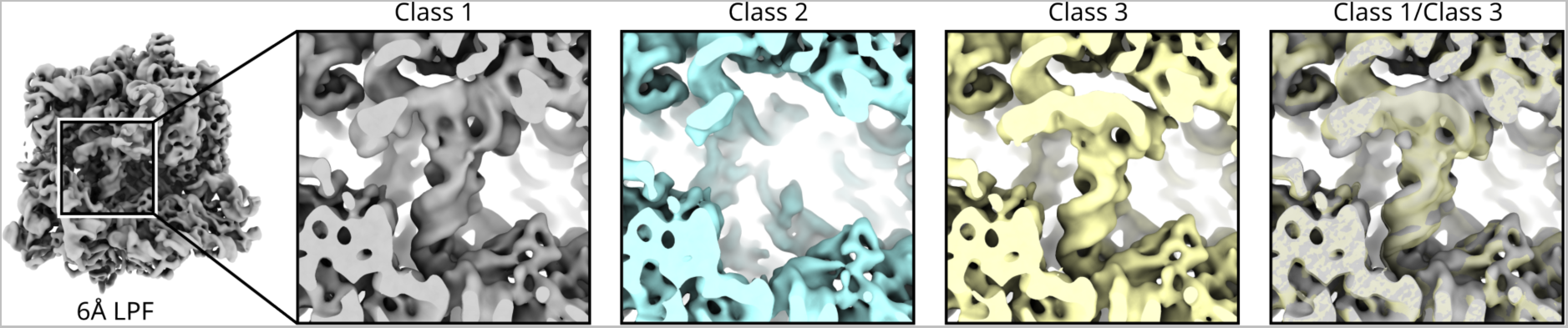
A substoichiometric tRNA occupies the A site of the VmlR2-70S complex. View of three sub-classified volumes (see **Supplementary Figure 3**) focusing on the A site. Volumes were low-pass-filtered to 6 Å. Inset shows A site with tRNA (classes 1 and 3) or without ligand (class 2). The right-most panel shows class 1 with transparent density for class 3 overlaid, revealing a different A-tRNA conformation.

**Supplementary Figure 7.**
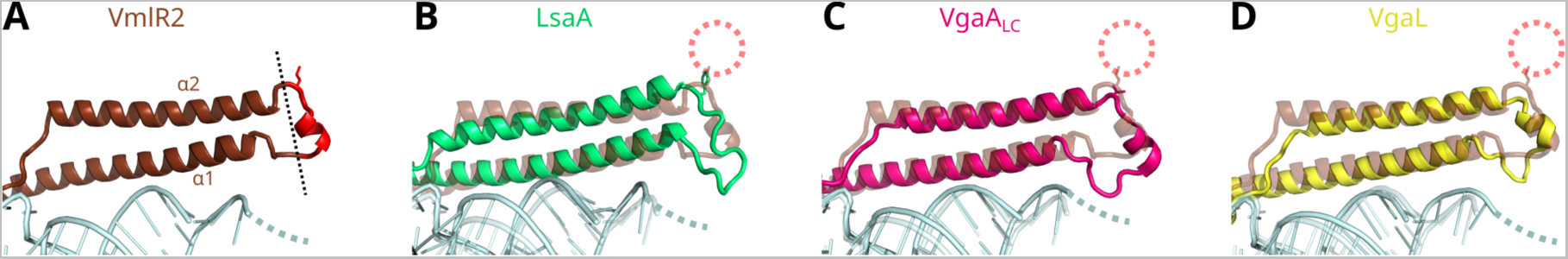
Comparison of interdomain linkers between selected ARE-ABCFs. (**A**) The VmlR2 interdomain linker (brown) with distorted P-tRNA (cyan). Dashed cyan lines indicate the approximate path of the tRNA CCA-3′ end, which was not modelled. Residues which are inserted compared to *B. subtilis* VmlR (see **Figure 1B**) are shown in red and demarcated by the black dotted line. Ile261, which reaches into the PTC, is shown as a stick representation. (**B**) Comparison of the LsaA (PDB ID 7NHK) and VmlR2 interdomain linkers. The red dashed circle indicates the approximate site of PLS_A_ binding. (**C**) As for **B** but with VgaA_LC_ (PDB 7NHL). (**D**) As for B but with VgaL (PDB ID 7NHN). Models were aligned by 23S rRNA.

**Supplementary Figure 8.**
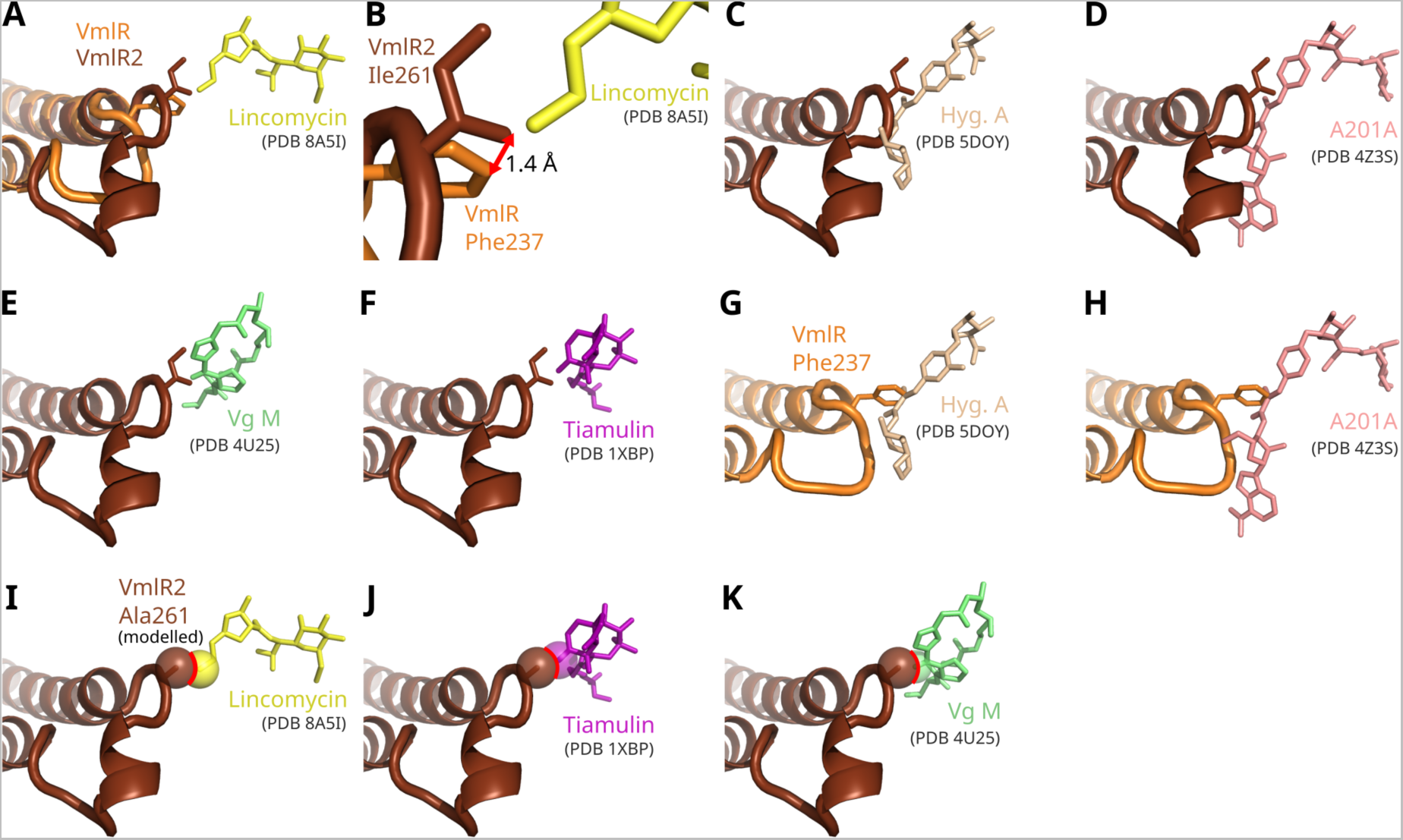
Overlap between ABCF-ARDs and selected antibiotics. (**A**) The VmlR2 (brown) and *B. subtilis* VmlR (orange) ARDs are similarly positioned with respect to the PLS_A_ binding site. For reference, lincomycin (PDB ID 8A5I) has been superimposed. (**B**) Close view of A, showing a 1.4 Å distance between the most-distal atoms of the VmlR2 and the *B. subtilis* VmlR ARDs. (**C**, **D**) Extensive predicted overlaps between the VmlR2 ARD and hygromycin A (**C**, beige, PDB ID 5DOY) and A201A (**D**, salmon, PDB ID 4Z3S) (Polikanov *et al*., 2015). (**E**, **F**) Same view as in **A** but with virginiamycin M (**E**, green, PDB ID 4U25) or tiamulin (**F**, purple, PDB ID 1XBP) superimposed (Schlunzen *et al*., 2004). (**G**, **H**) As for **C** and **D** but with the *B. subtilis* VmlR ARD still showing extensive overlaps. (**I-K**) Predicted overlaps between the VmlR2 I261A variant and lincomycin (**I**), tiamulin (**J**), and virginiamycin M (**K**) (Noeske et al., 2014). Transparent spheres represent van der Waals radii; red lines indicate clashes. Models were aligned by 23S rRNA.

**Supplementary Figure 9.**
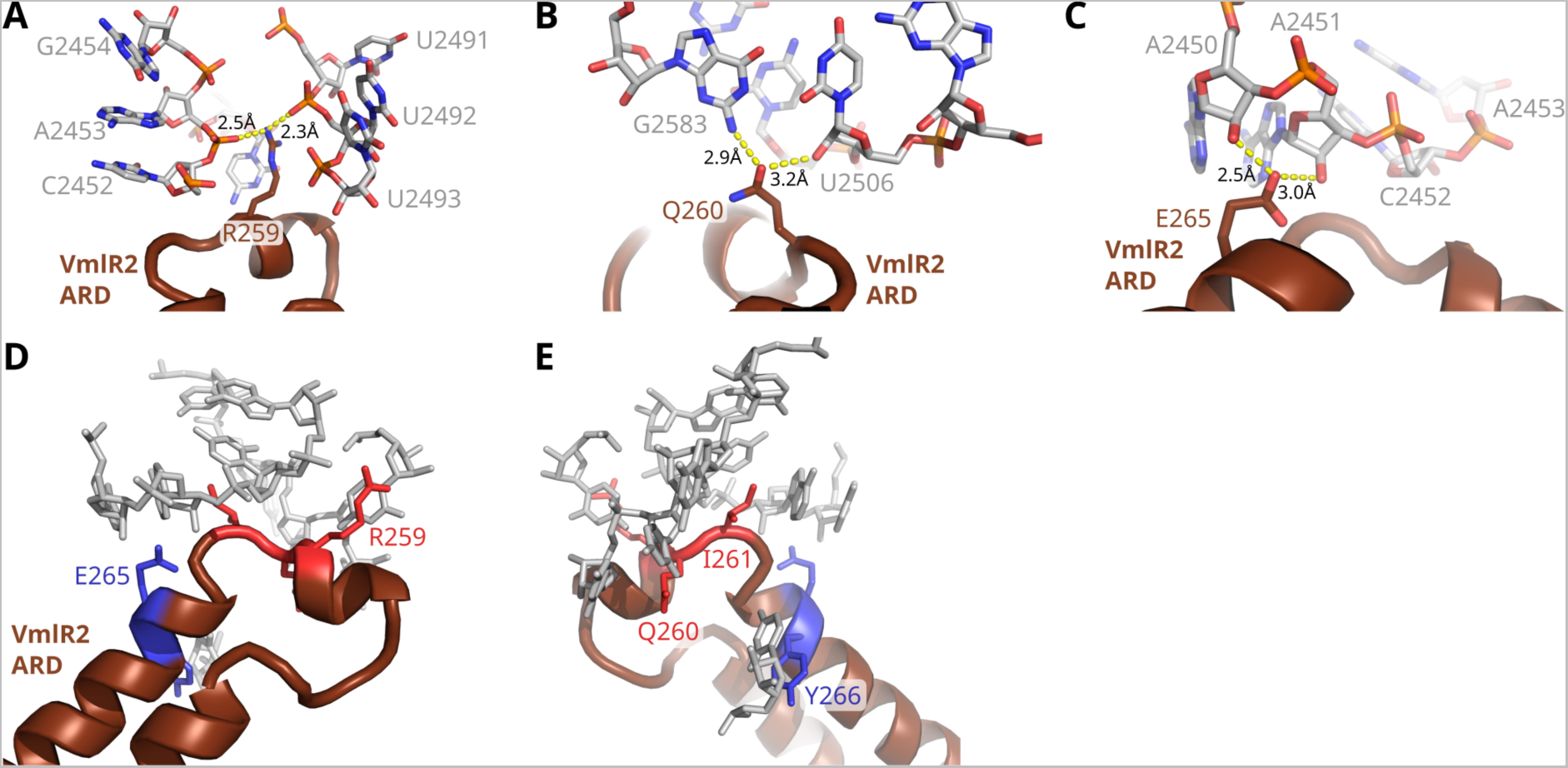
Interactions between the VmlR2 ARD and the 23S rRNA. (**A**) VmlR2 (brown) Arg259 forms H-bonds with 23S rRNA (grey) phosphate groups. (**B**) VmlR2 Gln260 interacts with G2583 and U2506 (**C**) VmlR2 E265 is within hydrogen-bonding distance of 23S rRNA A2450 and A2451. (**D**) Overview of alanine mutants tested in the VmlR2 ARD. Residues that, when substituted with alanine, substantially shifted the antibiotic resistance profile are shown in blue. Residues for which an alanine substitution had no effect are shown in red. (**E**) same as D but with a different view.

**Supplementary Figure 10.**
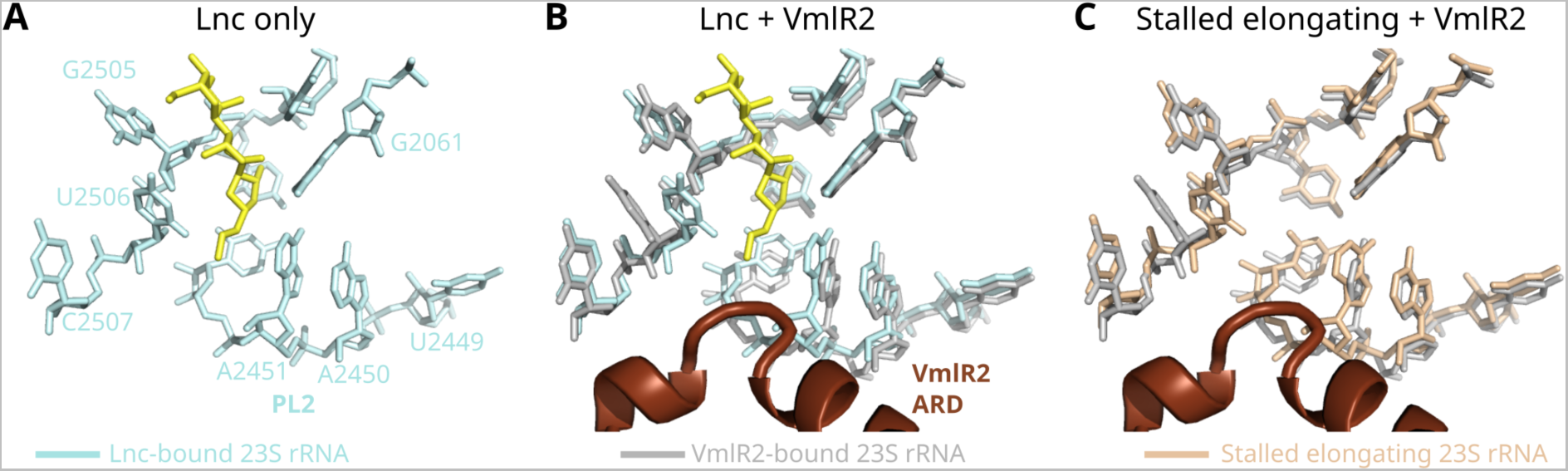
VmlR2 modulates the conformation of 23S rRNA nucleotides around the PLS_A_ binding site. (**A**) View of lincomycin binding with selected 23S rRNA nucleotides shown in cyan (PDB ID 8A5I). (**B**) Same as **A** but with the VmlR2 structure superimposed. 23S rRNA from the VmlR2 model is shown in grey. (**C**) Same as B, except the VmlR2 structure is compared to a stalled elongating *B. subtilis* 70S ribosome (PDB ID 6HA1).

**Supplementary Figure 11.**
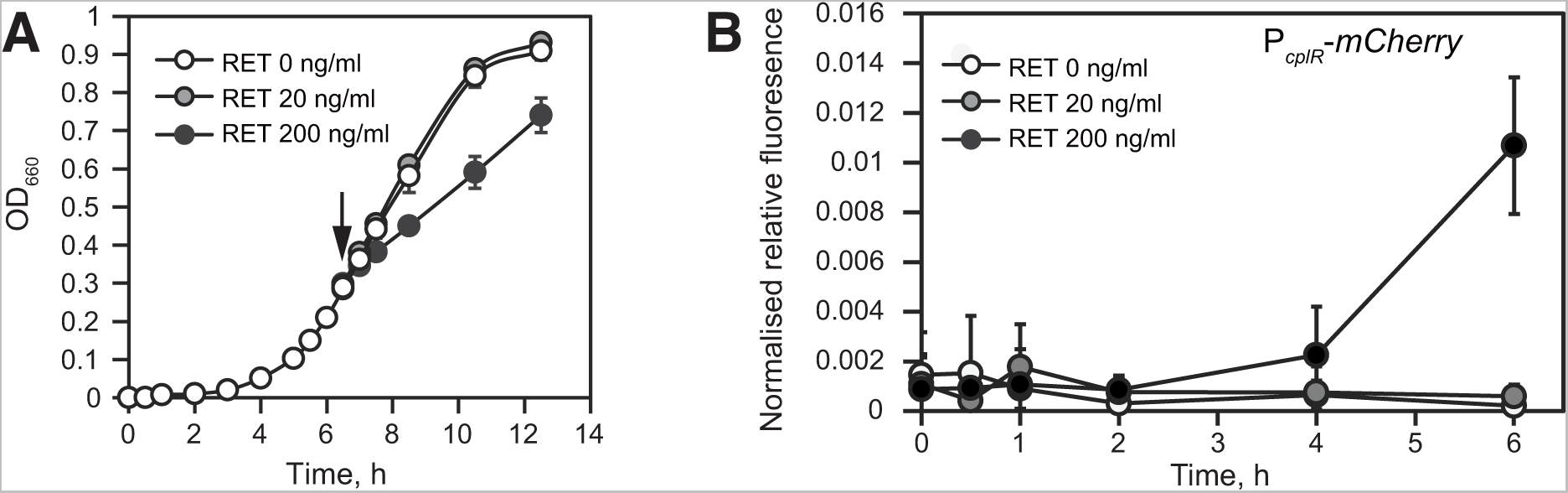
Expression of P*_cplR_*-*mCherry* fluorescent reporter is induced by retapamulin in a concentration-dependent manner. (**A**) Growth kinetics of *C. difficile* in the absence or presence of sub-MIC concentrations of retapamulin (20 and 200 ng/ml). (**B**) Kinetics of the induction of the P*_cplR_*-*mCherry* fluorescent reporter upon addition of retapamulin (RET) to final concentration of either 20 or 200 ng/ml. To calculate the relative promoter activity, relative fluorescence intensity normalised by O.D._600_ was further normalised by the signal from the constitutive P*_fdx_-mCherry* reporter.

## REFERENCES

Antimicrobial Resistance, C. (2022). Global burden of bacterial antimicrobial resistance in 2019: a systematic analysis. Lancet 399, 629–655. 10.1016/S0140-6736(21)02724-0.

Antonelli, A., D’Andrea, M.M., Brenciani, A., Galeotti, C.L., Morroni, G., Pollini, S., Varaldo, P.E., and Rossolini, G.M. (2018). Characterization of poxtA, a novel phenicol-oxazolidinone-tetracycline resistance gene from an MRSA of clinical origin. J Antimicrob Chemother 73, 1763–1769. 10.1093/jac/dky088.

Arenz, S., and Wilson, D.N. (2016). Bacterial Protein Synthesis as a Target for Antibiotic Inhibition. Cold Spring Harb Perspect Med 6. 10.1101/cshperspect.a025361.

Bannam, T.L., and Rood, J.I. (1993). Clostridium perfringens-Escherichia coli shuttle vectors that carry single antibiotic resistance determinants. Plasmid 29, 233–235. 10.1006/plas.1993.1025.

Barrasa, M.I., Tercero, J.A., Lacalle, R.A., and Jimenez, A. (1995). The ard1 gene from Streptomyces capreolus encodes a polypeptide of the ABC-transporters superfamily which confers resistance to the aminonucleoside antibiotic A201A. Eur J Biochem 228, 562–569. 10.1111/j.1432-1033.1995.tb20295.x.

Beckert, B., Leroy, E.C., Sothiselvam, S., Bock, L.V., Svetlov, M.S., Graf, M., Arenz, S., Abdelshahid, M., Seip, B., Grubmuller, H., et al. (2021). Structural and mechanistic basis for translation inhibition by macrolide and ketolide antibiotics. Nat Commun 12, 4466. 10.1038/s41467-021-24674-9.

Bishara, J., Peled, N., Pitlik, S., and Samra, Z. (2008). Mortality of patients with antibiotic-associated diarrhoea: the impact of Clostridium difficile. J Hosp Infect 68, 308–314. 10.1016/j.jhin.2008.01.033.

Boël, G., Smith, P.C., Ning, W., Englander, M.T., Chen, B., Hashem, Y., Testa, A.J., Fischer, J.J., Wieden, H.J., Frank, J., et al. (2014). The ABC-F protein EttA gates ribosome entry into the translation elongation cycle. Nat Struct Mol Biol 21, 143–151. 10.1038/nsmb.2740.

Britton, R.A., Eichenberger, P., Gonzalez-Pastor, J.E., Fawcett, P., Monson, R., Losick, R., and Grossman, A.D. (2002). Genome-wide analysis of the stationary-phase sigma factor (sigma-H) regulon of Bacillus subtilis. J Bacteriol 184, 4881–4890. 10.1128/JB.184.17.4881-4890.2002.

Brodiazhenko, T., Turnbull, K.J., Wu, K.J.Y., Takada, H., Tresco, B.I.C., Tenson, T., Myers, A.G., and Hauryliuk, V. (2022). Synthetic oxepanoprolinamide iboxamycin is active against Listeria monocytogenes despite the intrinsic resistance mediated by VgaL/Lmo0919 ABCF ATPase. JAC Antimicrob Resist 4, dlac061. 10.1093/jacamr/dlac061.

Buffie, C.G., Jarchum, I., Equinda, M., Lipuma, L., Gobourne, A., Viale, A., Ubeda, C., Xavier, J., and Pamer, E.G. (2012). Profound alterations of intestinal microbiota following a single dose of clindamycin results in sustained susceptibility to Clostridium difficile-induced colitis. Infect Immun 80, 62–73. 10.1128/IAI.05496-11.

Cai, X., Zhan, Y., Cao, Z., Yan, B., and Cai, J. (2021). Expression of ribosomal protection protein RppA is regulated by a ribosome-dependent ribo-regulator and two mistranslation products. Environ Microbiol 23, 696–712. 10.1111/1462-2920.15143.

Camacho, C., Coulouris, G., Avagyan, V., Ma, N., Papadopoulos, J., Bealer, K., and Madden, T.L. (2009). BLAST+: architecture and applications. BMC Bioinformatics 10, 421. 10.1186/1471-2105-10-421.

Candiano, G., Bruschi, M., Musante, L., Santucci, L., Ghiggeri, G.M., Carnemolla, B., Orecchia, P., Zardi, L., and Righetti, P.G. (2004). Blue silver: a very sensitive colloidal Coomassie G-250 staining for proteome analysis. Electrophoresis 25, 1327–1333. 10.1002/elps.200305844.

Capella-Gutierrez, S., Silla-Martinez, J.M., and Gabaldon, T. (2009). trimAl: a tool for automated alignment trimming in large-scale phylogenetic analyses. Bioinformatics 25, 1972–1973. 10.1093/bioinformatics/btp348.

Casanal, A., Lohkamp, B., and Emsley, P. (2020). Current developments in Coot for macromolecular model building of Electron Cryo-microscopy and Crystallographic Data. Protein Sci 29, 1069–1078. 10.1002/pro.3791.

Chen, B., Boël, G., Hashem, Y., Ning, W., Fei, J., Wang, C., Gonzalez, R.L., Jr., Hunt, J.F., and Frank, J. (2014). EttA regulates translation by binding the ribosomal E site and restricting ribosome-tRNA dynamics. Nat Struct Mol Biol 21, 152–159. 10.1038/nsmb.2741.

Crowe-McAuliffe, C., Graf, M., Huter, P., Takada, H., Abdelshahid, M., Novacek, J., Murina, V., Atkinson, G.C., Hauryliuk, V., and Wilson, D.N. (2018). Structural basis for antibiotic resistance mediated by the Bacillus subtilis ABCF ATPase VmlR. Proc Natl Acad Sci U S A 115, 8978–8983. 10.1073/pnas.1808535115.

Crowe-McAuliffe, C., Murina, V., Turnbull, K.J., Huch, S., Kasari, M., Takada, H., Nersisyan, L., Sundsfjord, A., Hegstad, K., Atkinson, G.C., et al. (2022). Structural basis for PoxtA-mediated resistance to phenicol and oxazolidinone antibiotics. Nat Commun 13, 1860. 10.1038/s41467-022-29274-9.

Crowe-McAuliffe, C., Murina, V., Turnbull, K.J., Kasari, M., Mohamad, M., Polte, C., Takada, H., Vaitkevicius, K., Johansson, J., Ignatova, Z., et al. (2021). Structural basis of ABCF-mediated resistance to pleuromutilin, lincosamide, and streptogramin A antibiotics in Gram-positive pathogens. Nat Commun 12, 3577. 10.1038/s41467-021-23753-1.

Cui, Z., Li, X., Shin, J., Gamper, H., Hou, Y.M., Sacchettini, J.C., and Zhang, J. (2022). Interplay between an ATP-binding cassette F protein and the ribosome from Mycobacterium tuberculosis. Nat Commun 13, 432. 10.1038/s41467-022-28078-1.

Czepiel, J., Drozdz, M., Pituch, H., Kuijper, E.J., Perucki, W., Mielimonka, A., Goldman, S., Wultanska, D., Garlicki, A., and Biesiada, G. (2019). Clostridium difficile infection: review. Eur J Clin Microbiol Infect Dis 38, 1211–1221. 10.1007/s10096-019-03539-6.

Dar, D., Shamir, M., Mellin, J.R., Koutero, M., Stern-Ginossar, N., Cossart, P., and Sorek, R. (2016). Term-seq reveals abundant ribo-regulation of antibiotics resistance in bacteria. Science 352, aad9822. 10.1126/science.aad9822.

Davis, A.R., Gohara, D.W., and Yap, M.N. (2014). Sequence selectivity of macrolide-induced translational attenuation. Proc Natl Acad Sci U S A 111, 15379–15384. 10.1073/pnas.1410356111.

Duffy, C.R., Huang, Y., Andrikopoulou, M., Stern-Ascher, C.N., Wright, J.D., Goffman, D., D’Alton, M.E., and Friedman, A.M. (2020). Clindamycin, Gentamicin, and Risk of Clostridium difficile Infection and Acute Kidney Injury During Delivery Hospitalizations. Obstet Gynecol 135, 59–67. 10.1097/AOG.0000000000003568.

Egorov, A.A., and Atkinson, G.C. (2022). uORF4u: a tool for annotation of conserved upstream open reading frames. bioRxiv, 2022.2010.2027.514069. 10.1101/2022.10.27.514069.

Egorov, A.A., Sakharova, E.A., Anisimova, A.S., Dmitriev, S.E., Gladyshev, V.N., and Kulakovskiy, I.V. (2019). svist4get: a simple visualization tool for genomic tracks from sequencing experiments. BMC Bioinformatics 20, 113. 10.1186/s12859-019-2706-8.

Ero, R., Kumar, V., Su, W., and Gao, Y.G. (2019). Ribosome protection by ABC-F proteins-Molecular mechanism and potential drug design. Protein Sci 28, 684–693. 10.1002/pro.3589.

Fagan, R.P., and Fairweather, N.F. (2011). Clostridium difficile has two parallel and essential Sec secretion systems. J Biol Chem 286, 27483–27493. 10.1074/jbc.M111.263889.

Farnham, P.J., and Platt, T. (1981). Rho-independent termination: dyad symmetry in DNA causes RNA polymerase to pause during transcription in vitro. Nucleic Acids Res 9, 563–577. 10.1093/nar/9.3.563.

Farrow, K.A., Lyras, D., and Rood, J.I. (2001). Genomic analysis of the erythromycin resistance element Tn5398 from Clostridium difficile. Microbiology (Reading) 147, 2717–2728. 10.1099/00221287-147-10-2717.

Fuchs, M., Lamm-Schmidt, V., Sulzer, J., Ponath, F., Jenniches, L., Kirk, J.A., Fagan, R.P., Barquist, L., Vogel, J., and Faber, F. (2021). An RNA-centric global view of Clostridioides difficile reveals broad activity of Hfq in a clinically important gram-positive bacterium. Proc Natl Acad Sci U S A 118. 10.1073/pnas.2103579118.

Gentry, D.R., McCloskey, L., Gwynn, M.N., Rittenhouse, S.F., Scangarella, N., Shawar, R., and Holmes, D.J. (2008). Genetic characterization of Vga ABC proteins conferring reduced susceptibility to pleuromutilins in Staphylococcus aureus. Antimicrob Agents Chemother 52, 4507–4509. 10.1128/AAC.00915-08.

Gruber, A.R., Lorenz, R., Bernhart, S.H., Neubock, R., and Hofacker, I.L. (2008). The Vienna RNA websuite. Nucleic Acids Res 36, W70–74. 10.1093/nar/gkn188.

Gumkowski, J.D., Martinie, R.J., Corrigan, P.S., Pan, J., Bauerle, M.R., Almarei, M., Booker, S.J., Silakov, A., Krebs, C., and Boal, A.K. (2019). Analysis of RNA Methylation by Phylogenetically Diverse Cfr Radical S-Adenosylmethionine Enzymes Reveals an Iron-Binding Accessory Domain in a Clostridial Enzyme. Biochemistry 58, 3169–3184. 10.1021/acs.biochem.9b00197.

Gusarov, I., and Nudler, E. (1999). The mechanism of intrinsic transcription termination. Mol Cell 3, 495–504. 10.1016/s1097-2765(00)80477-3.

Harms, J.M., Schlunzen, F., Fucini, P., Bartels, H., and Yonath, A. (2004). Alterations at the peptidyl transferase centre of the ribosome induced by the synergistic action of the streptogramins dalfopristin and quinupristin. BMC Biol 2, 4. 10.1186/1741-7007-2-4.

Ippolito, J.A., Kanyo, Z.F., Wang, D., Franceschi, F.J., Moore, P.B., Steitz, T.A., and Duffy, E.M. (2008). Crystal structure of the oxazolidinone antibiotic linezolid bound to the 50S ribosomal subunit. J Med Chem 51, 3353–3356. 10.1021/jm800379d.

Jamali, K., Kimanius, D., and Scheres, S. (2022). ModelAngelo: Automated Model Building in Cryo-EM Maps. arXiv preprint arXiv:2210.00006.

Johnson, S., Samore, M.H., Farrow, K.A., Killgore, G.E., Tenover, F.C., Lyras, D., Rood, J.I., DeGirolami, P., Baltch, A.L., Rafferty, M.E., et al. (1999). Epidemics of diarrhea caused by a clindamycin-resistant strain of Clostridium difficile in four hospitals. N Engl J Med 341, 1645–1651. 10.1056/NEJM199911253412203.

Kaminishi, T., Schedlbauer, A., Fabbretti, A., Brandi, L., Ochoa-Lizarralde, B., He, C.G., Milon, P., Connell, S.R., Gualerzi, C.O., and Fucini, P. (2015). Crystallographic characterization of the ribosomal binding site and molecular mechanism of action of Hygromycin A. Nucleic Acids Res 43, 10015–10025. 10.1093/nar/gkv975.

Kannan, K., Kanabar, P., Schryer, D., Florin, T., Oh, E., Bahroos, N., Tenson, T., Weissman, J.S., and Mankin, A.S. (2014). The general mode of translation inhibition by macrolide antibiotics. Proc Natl Acad Sci U S A 111, 15958–15963. 10.1073/pnas.1417334111.

Katoh, K., and Standley, D.M. (2013). MAFFT multiple sequence alignment software version 7: improvements in performance and usability. Mol Biol Evol 30, 772–780. 10.1093/molbev/mst010.

Kehrenberg, C., Aarestrup, F.M., and Schwarz, S. (2007). IS21-558 insertion sequences are involved in the mobility of the multiresistance gene cfr. Antimicrob Agents Chemother 51, 483–487. 10.1128/AAC.01340-06.

Koberska, M., Vesela, L., Vimberg, V., Lenart, J., Vesela, J., Kamenik, Z., Janata, J., and Balikova Novotna, G. (2021). Beyond Self-Resistance: ABCF ATPase LmrC Is a Signal-Transducing Component of an Antibiotic-Driven Signaling Cascade Accelerating the Onset of Lincomycin Biosynthesis. mBio 12, e0173121. 10.1128/mBio.01731-21.

Koller, T.O., Turnbull, K.J., Vaitkevicius, K., Crowe-McAuliffe, C., Roghanian, M., Bulvas, O., Nakamoto, J.A., Kurata, T., Julius, C., Atkinson, G.C., et al. (2022). Structural basis for HflXr-mediated antibiotic resistance in Listeria monocytogenes. Nucleic Acids Res 50, 11285–11300. 10.1093/nar/gkac934.

Kovalevskiy, O., Nicholls, R.A., Long, F., Carlon, A., and Murshudov, G.N. (2018). Overview of refinement procedures within REFMAC5: utilizing data from different sources. Acta Crystallogr D Struct Biol 74, 215–227. 10.1107/S2059798318000979.

Lai, C.J., and Weisblum, B. (1971). Altered methylation of ribosomal RNA in an erythromycin-resistant strain of Staphylococcus aureus. Proc Natl Acad Sci U S A 68, 856–860. 10.1073/pnas.68.4.856.

Larsson, A. (2014). AliView: a fast and lightweight alignment viewer and editor for large datasets. Bioinformatics 30, 3276–3278. 10.1093/bioinformatics/btu531.

Leclercq, R., and Courvalin, P. (1991). Bacterial resistance to macrolide, lincosamide, and streptogramin antibiotics by target modification. Antimicrob Agents Chemother 35, 1267–1272. 10.1128/AAC.35.7.1267.

Liebschner, D., Afonine, P.V., Baker, M.L., Bunkoczi, G., Chen, V.B., Croll, T.I., Hintze, B., Hung, L.W., Jain, S., McCoy, A.J., et al. (2019). Macromolecular structure determination using X-rays, neutrons and electrons: recent developments in Phenix. Acta Crystallogr D Struct Biol 75, 861–877. 10.1107/S2059798319011471.

Lin, J., Zhou, D., Steitz, T.A., Polikanov, Y.S., and Gagnon, M.G. (2018). Ribosome-Targeting Antibiotics: Modes of Action, Mechanisms of Resistance, and Implications for Drug Design. Annu Rev Biochem 87, 451–478. 10.1146/annurev-biochem-062917-011942.

Long, K.S., Poehlsgaard, J., Kehrenberg, C., Schwarz, S., and Vester, B. (2006). The Cfr rRNA methyltransferase confers resistance to Phenicols, Lincosamides, Oxazolidinones, Pleuromutilins, and Streptogramin A antibiotics. Antimicrob Agents Chemother 50, 2500–2505. 10.1128/AAC.00131-06.

Ma, N.J., Moonan, D.W., and Isaacs, F.J. (2014). Precise manipulation of bacterial chromosomes by conjugative assembly genome engineering. Nat Protoc 9, 2285–2300. 10.1038/nprot.2014.081.

Macleod, A.J., Ross, H.B., Ozere, R.L., Digout, G., and Van, R. (1964). Lincomycin: A New Antibiotic Active against Staphylococci and Other Gram-Positive Cocci: Clinical and Laboratory Studies. Can Med Assoc J 91, 1056–1060.

Maravić, G. (2004). Macrolide resistance based on the Erm-mediated rRNA methylation. Curr Drug Targets Infect Disord 4, 193–202. 10.2174/1568005043340777.

Martin, J.S., Monaghan, T.M., and Wilcox, M.H. (2016). Clostridium difficile infection: epidemiology, diagnosis and understanding transmission. Nat Rev Gastroenterol Hepatol 13, 206–216. 10.1038/nrgastro.2016.25.

Miller, M.A., Pfeiffer, W., and Schwartz, T. (2010). Creating the CIPRES Science Gateway for inference of large phylogenetic trees. 14-14 Nov. 2010. pp. 1–8.

Minh, B.Q., Schmidt, H.A., Chernomor, O., Schrempf, D., Woodhams, M.D., von Haeseler, A., and Lanfear, R. (2020). IQ-TREE 2: New Models and Efficient Methods for Phylogenetic Inference in the Genomic Era. Mol Biol Evol 37, 1530–1534. 10.1093/molbev/msaa015.

Mitcheltree, M.J., Pisipati, A., Syroegin, E.A., Silvestre, K.J., Klepacki, D., Mason, J.D., Terwilliger, D.W., Testolin, G., Pote, A.R., Wu, K.J.Y., et al. (2021). A synthetic antibiotic class overcoming bacterial multidrug resistance. Nature 599, 507–512. 10.1038/s41586-021-04045-6.

Mohamad, M., Nicholson, D., Saha, C.K., Hauryliuk, V., Edwards, T.A., Atkinson, G.C., Ranson, N.A., and O’Neill, A.J. (2022). Sal-type ABC-F proteins: intrinsic and common mediators of pleuromutilin resistance by target protection in staphylococci. Nucleic Acids Res 50, 2128–2142. 10.1093/nar/gkac058.

Murina, V., Kasari, M., Hauryliuk, V., and Atkinson, G.C. (2018). Antibiotic resistance ABCF proteins reset the peptidyl transferase centre of the ribosome to counter translational arrest. Nucleic Acids Res 46, 3753–3763. 10.1093/nar/gky050.

Murina, V., Kasari, M., Takada, H., Hinnu, M., Saha, C.K., Grimshaw, J.W., Seki, T., Reith, M., Putrins, M., Tenson, T., et al. (2019). ABCF ATPases Involved in Protein Synthesis, Ribosome Assembly and Antibiotic Resistance: Structural and Functional Diversification across the Tree of Life. J Mol Biol 431, 3568–3590. 10.1016/j.jmb.2018.12.013.

Nariya, H., Miyata, S., Suzuki, M., Tamai, E., and Okabe, A. (2011). Development and application of a method for counterselectable in-frame deletion in Clostridium perfringens. Appl Environ Microbiol 77, 1375–1382. 10.1128/AEM.01572-10.

Noeske, J., Huang, J., Olivier, N.B., Giacobbe, R.A., Zambrowski, M., and Cate, J.H. (2014). Synergy of streptogramin antibiotics occurs independently of their effects on translation. Antimicrob Agents Chemother 58, 5269–5279. 10.1128/AAC.03389-14.

Novotna, G., and Janata, J. (2006). A new evolutionary variant of the streptogramin A resistance protein, Vga(A)LC, from Staphylococcus haemolyticus with shifted substrate specificity towards lincosamides. Antimicrob Agents Chemother 50, 4070–4076. 10.1128/AAC.00799-06.

Obana, N., Nakamura, K., and Nomura, N. (2020). Temperature-regulated heterogeneous extracellular matrix gene expression defines biofilm morphology in Clostridium perfringens. NPJ Biofilms Microbiomes 6, 29. 10.1038/s41522-020-00139-7.

Obana, N., Shirahama, Y., Abe, K., and Nakamura, K. (2010). Stabilization of Clostridium perfringens collagenase mRNA by VR-RNA-dependent cleavage in 5’ leader sequence. Mol Microbiol 77, 1416–1428. 10.1111/j.1365-2958.2010.07258.x.

Ohki, R., Tateno, K., Takizawa, T., Aiso, T., and Murata, M. (2005). Transcriptional termination control of a novel ABC transporter gene involved in antibiotic resistance in Bacillus subtilis. J Bacteriol 187, 5946–5954. 10.1128/JB.187.17.5946-5954.2005.

Orelle, C., Carlson, S., Kaushal, B., Almutairi, M.M., Liu, H., Ochabowicz, A., Quan, S., Pham, V.C., Squires, C.L., Murphy, B.T., and Mankin, A.S. (2013). Tools for characterizing bacterial protein synthesis inhibitors. Antimicrob Agents Chemother 57, 5994–6004. 10.1128/AAC.01673-13.

Peltier, J., Hamiot, A., Garneau, J.R., Boudry, P., Maikova, A., Hajnsdorf, E., Fortier, L.C., Dupuy, B., and Soutourina, O. (2020). Type I toxin-antitoxin systems contribute to the maintenance of mobile genetic elements in Clostridioides difficile. Commun Biol 3, 718. 10.1038/s42003-020-01448-5.

Pettersen, E.F., Goddard, T.D., Huang, C.C., Meng, E.C., Couch, G.S., Croll, T.I., Morris, J.H., and Ferrin, T.E. (2021). UCSF ChimeraX: Structure visualization for researchers, educators, and developers. Protein Sci 30, 70–82. 10.1002/pro.3943.

Phillips, I. (1981). Past and current use of clindamycin and lincomycin. J Antimicrob Chemother 7 Suppl A, 11–18. 10.1093/jac/7.suppl_a.11.

Polikanov, Y.S., Starosta, A.L., Juette, M.F., Altman, R.B., Terry, D.S., Lu, W., Burnett, B.J., Dinos, G., Reynolds, K.A., Blanchard, S.C., et al. (2015). Distinct tRNA Accommodation Intermediates Observed on the Ribosome with the Antibiotics Hygromycin A and A201A. Mol Cell 58, 832–844. 10.1016/j.molcel.2015.04.014.

Polikanov, Y.S., Steitz, T.A., and Innis, C.A. (2014). A proton wire to couple aminoacyl-tRNA accommodation and peptide-bond formation on the ribosome. Nat Struct Mol Biol 21, 787–793. 10.1038/nsmb.2871.

Ransom, E.M., Ellermeier, C.D., and Weiss, D.S. (2015). Use of mCherry Red fluorescent protein for studies of protein localization and gene expression in Clostridium difficile. Appl Environ Microbiol 81, 1652–1660. 10.1128/AEM.03446-14.

Roberts, A.P., and Smits, W.K. (2018). The evolving epidemic of Clostridium difficile 630. Anaerobe 53, 2–4. 10.1016/j.anaerobe.2018.04.015.

Ross, J.I., Eady, E.A., Cove, J.H., Cunliffe, W.J., Baumberg, S., and Wootton, J.C. (1990). Inducible erythromycin resistance in staphylococci is encoded by a member of the ATP-binding transport super-gene family. Mol Microbiol 4, 1207–1214. 10.1111/j.1365-2958.1990.tb00696.x.

Rosteck, P.R., Jr., Reynolds, P.A., and Hershberger, C.L. (1991). Homology between proteins controlling Streptomyces fradiae tylosin resistance and ATP-binding transport. Gene 102, 27–32. 10.1016/0378-1119(91)90533-h.

Saugar, I., Sanz, E., Rubio, M.A., Espinosa, J.C., and Jimenez, A. (2002). Identification of a set of genes involved in the biosynthesis of the aminonucleoside moiety of antibiotic A201A from Streptomyces capreolus. Eur J Biochem 269, 5527–5535. 10.1046/j.1432-1033.2002.03258.x.

Scheres, S.H. (2012). RELION: implementation of a Bayesian approach to cryo-EM structure determination. J Struct Biol 180, 519–530. 10.1016/j.jsb.2012.09.006.

Schlunzen, F., Pyetan, E., Fucini, P., Yonath, A., and Harms, J.M. (2004). Inhibition of peptide bond formation by pleuromutilins: the structure of the 50S ribosomal subunit from Deinococcus radiodurans in complex with tiamulin. Mol Microbiol 54, 1287–1294. 10.1111/j.1365-2958.2004.04346.x.

Schwarz, S., Shen, J., Kadlec, K., Wang, Y., Brenner Michael, G., Fessler, A.T., and Vester, B. (2016). Lincosamides, Streptogramins, Phenicols, and Pleuromutilins: Mode of Action and Mechanisms of Resistance. Cold Spring Harb Perspect Med 6. 10.1101/cshperspect.a027037.

Schwarz, S., Werckenthin, C., and Kehrenberg, C. (2000). Identification of a plasmid-borne chloramphenicol-florfenicol resistance gene in Staphylococcus sciuri. Antimicrob Agents Chemother 44, 2530–2533. 10.1128/AAC.44.9.2530-2533.2000.

Seip, B., Sacheau, G., Dupuy, D., and Innis, C.A. (2018). Ribosomal stalling landscapes revealed by high-throughput inverse toeprinting of mRNA libraries. Life Sci Alliance 1, e201800148. 10.26508/lsa.201800148.

Singh, K.V., Weinstock, G.M., and Murray, B.E. (2002). An Enterococcus faecalis ABC homologue (Lsa) is required for the resistance of this species to clindamycin and quinupristin-dalfopristin. Antimicrob Agents Chemother 46, 1845–1850. 10.1128/aac.46.6.1845-1850.2002.

Sipos, K., Szigeti, R., Dong, X., and Turnbough, C.L., Jr. (2007). Systematic mutagenesis of the thymidine tract of the pyrBI attenuator and its effects on intrinsic transcription termination in Escherichia coli. Mol Microbiol 66, 127–138. 10.1111/j.1365-2958.2007.05902.x.

Stabler, R.A., He, M., Dawson, L., Martin, M., Valiente, E., Corton, C., Lawley, T.D., Sebaihia, M., Quail, M.A., Rose, G., et al. (2009). Comparative genome and phenotypic analysis of Clostridium difficile 027 strains provides insight into the evolution of a hypervirulent bacterium. Genome Biol 10, R102. 10.1186/gb-2009-10-9-r102.

Su, W., Kumar, V., Ding, Y., Ero, R., Serra, A., Lee, B.S.T., Wong, A.S.W., Shi, J., Sze, S.K., Yang, L., and Gao, Y.G. (2018). Ribosome protection by antibiotic resistance ATP-binding cassette protein. Proc Natl Acad Sci U S A 115, 5157–5162. 10.1073/pnas.1803313115.

Sullivan, A., Edlund, C., and Nord, C.E. (2001). Effect of antimicrobial agents on the ecological balance of human microflora. Lancet Infect Dis 1, 101–114. 10.1016/S1473-3099(01)00066-4.

Syroegin, E.A., Aleksandrova, E.V., and Polikanov, Y.S. (2022a). Structural basis for the inability of chloramphenicol to inhibit peptide bond formation in the presence of A-site glycine. Nucleic Acids Res 50, 7669–7679. 10.1093/nar/gkac548.

Syroegin, E.A., Flemmich, L., Klepacki, D., Vazquez-Laslop, N., Micura, R., and Polikanov, Y.S. (2022b). Structural basis for the context-specific action of the classic peptidyl transferase inhibitor chloramphenicol. Nat Struct Mol Biol 29, 152–161. 10.1038/s41594-022-00720-y.

Takada, H., Mandell, Z.F., Yakhnin, H., Glazyrina, A., Chiba, S., Kurata, T., Wu, K.J.Y., Tresco, B.I.C., Myers, A.G., Aktinson, G.C., et al. (2022). Expression of Bacillus subtilis ABCF antibiotic resistance factor VmlR is regulated by RNA polymerase pausing, transcription attenuation, translation attenuation and (p)ppGpp. Nucleic Acids Res. 10.1093/nar/gkac497.

Takamizawa, A., Miyata, S., Matsushita, O., Kaji, M., Taniguchi, Y., Tamai, E., Shimamoto, S., and Okabe, A. (2004). High-level expression of clostridial sialidase using a ferredoxin gene promoter-based plasmid. Protein Expr Purif 36, 70–75. 10.1016/j.pep.2004.03.004.

Tareen, A., and Kinney, J.B. (2020). Logomaker: beautiful sequence logos in Python. Bioinformatics 36, 2272–2274. 10.1093/bioinformatics/btz921.

Tsai, K., Stojkovic, V., Lee, D.J., Young, I.D., Szal, T., Klepacki, D., Vazquez-Laslop, N., Mankin, A.S., Fraser, J.S., and Fujimori, D.G. (2022). Structural basis for context-specific inhibition of translation by oxazolidinone antibiotics. Nat Struct Mol Biol 29, 162–171. 10.1038/s41594-022-00723-9.

Tu, D., Blaha, G., Moore, P.B., and Steitz, T.A. (2005). Structures of MLSBK antibiotics bound to mutated large ribosomal subunits provide a structural explanation for resistance. Cell 121, 257–270. 10.1016/j.cell.2005.02.005.

Uchiyama, H., and Weisblum, B. (1985). N-Methyl transferase of Streptomyces erythraeus that confers resistance to the macrolide-lincosamide-streptogramin B antibiotics: amino acid sequence and its homology to cognate R-factor enzymes from pathogenic bacilli and cocci. Gene 38, 103–110. 10.1016/0378-1119(85)90208-2.

Varadi, M., Anyango, S., Deshpande, M., Nair, S., Natassia, C., Yordanova, G., Yuan, D., Stroe, O., Wood, G., Laydon, A., et al. (2022). AlphaFold Protein Structure Database: massively expanding the structural coverage of protein-sequence space with high-accuracy models. Nucleic Acids Res 50, D439–D444. 10.1093/nar/gkab1061.

Vazquez-Laslop, N., and Mankin, A.S. (2018). Context-Specific Action of Ribosomal Antibiotics Annu Rev Microbiol 72, 185–207. 10.1146/annurev-micro-090817-062329.

Vimberg, V., Cavanagh, J.P., Novotna, M., Lenart, J., Nguyen Thi Ngoc, B., Vesela, J., Pain, M., Koberska, M., and Balikova Novotna, G. (2020). Ribosome-Mediated Attenuation of vga(A) Expression Is Shaped by the Antibiotic Resistance Specificity of Vga(A) Protein Variants. Antimicrob Agents Chemother 64. 10.1128/AAC.00666-20.

Wang, Y., Lv, Y., Cai, J., Schwarz, S., Cui, L., Hu, Z., Zhang, R., Li, J., Zhao, Q., He, T., et al. (2015). A novel gene, optrA, that confers transferable resistance to oxazolidinones and phenicols and its presence in Enterococcus faecalis and Enterococcus faecium of human and animal origin. J Antimicrob Chemother 70, 2182–2190. 10.1093/jac/dkv116.

Waterhouse, A.M., Procter, J.B., Martin, D.M., Clamp, M., and Barton, G.J. (2009). Jalview Version 2--a multiple sequence alignment editor and analysis workbench. Bioinformatics 25, 1189–1191. 10.1093/bioinformatics/btp033.

Watson, Z.L., Ward, F.R., Meheust, R., Ad, O., Schepartz, A., Banfield, J.F., and Cate, J.H. (2020). Structure of the bacterial ribosome at 2 A resolution. Elife 9. 10.7554/eLife.60482.

Williams, C.J., Headd, J.J., Moriarty, N.W., Prisant, M.G., Videau, L.L., Deis, L.N., Verma, V., Keedy, D.A., Hintze, B.J., Chen, V.B., et al. (2018). MolProbity: More and better reference data for improved all-atom structure validation. Protein Sci 27, 293–315. 10.1002/pro.3330.

Wilson, D.N. (2014). Ribosome-targeting antibiotics and mechanisms of bacterial resistance. Nat Rev Microbiol 12, 35–48. 10.1038/nrmicro3155.

Wilson, D.N., Hauryliuk, V., Atkinson, G.C., and O’Neill, A.J. (2020). Target protection as a key antibiotic resistance mechanism. Nat Rev Microbiol 18, 637–648. 10.1038/s41579-020-0386-z.

Yamashita, K., Palmer, C.M., Burnley, T., and Murshudov, G.N. (2021). Cryo-EM single-particle structure refinement and map calculation using Servalcat. Acta Crystallogr D Struct Biol 77, 1282–1291. 10.1107/S2059798321009475.

Yarnell, W.S., and Roberts, J.W. (1999). Mechanism of intrinsic transcription termination and antitermination. Science 284, 611–615. 10.1126/science.284.5414.611.

Zhang, K. (2016). Gctf: Real-time CTF determination and correction. J Struct Biol 193, 1–12. 10.1016/j.jsb.2015.11.003.

Zheng, S.Q., Palovcak, E., Armache, J.P., Verba, K.A., Cheng, Y., and Agard, D.A. (2017). MotionCor2: anisotropic correction of beam-induced motion for improved cryo-electron microscopy. Nat Methods 14, 331–332. 10.1038/nmeth.4193.

Zhou, L., Feng, T., Xu, S., Gao, F., Lam, T.T., Wang, Q., Wu, T., Huang, H., Zhan, L., Li, L., et al. (2022). ggmsa: a visual exploration tool for multiple sequence alignment and associated data. Brief Bioinform 23. 10.1093/bib/bbac222.

Zivanov, J., Nakane, T., Forsberg, B.O., Kimanius, D., Hagen, W.J., Lindahl, E., and Scheres, S.H. (2018). New tools for automated high-resolution cryo-EM structure determination in RELION-3. Elife 7. 10.7554/eLife.42166.

Zuker, M. (2003). Mfold web server for nucleic acid folding and hybridization prediction. Nucleic Acids Res 31, 3406–3415. 10.1093/nar/gkg595.

